# Multistability in cellular differentiation enabled by a network of three mutually repressing master regulators

**DOI:** 10.1101/2020.05.14.089805

**Authors:** Atchuta Srinivas Duddu, Sarthak Sahoo, Souvadra Hati, Siddharth Jhunjhunwala, Mohit Kumar Jolly

**Author notes:** Author to whom correspondence to be addressed (M.K.J.).

## Abstract

Identifying the design principles of complex regulatory networks driving cellular decision-making remains essential to decode embryonic development as well as enhance cellular reprogramming. A well-studied network motif involved in cellular decision-making is a toggle switch – a set of two opposing transcription factors A and B, each of which is a master regulator of a specific cell-fate and can inhibit the activity of the other. A toggle switch can lead to two possible states – (high A, low B) and (low A, high B), and drives the ‘either-or’ choice between these two cell-fates for a common progenitor cell. However, the principles of coupled toggle switches remains unclear. Here, we investigate the dynamics of three master regulators A, B and C inhibiting each other, thus forming three coupled toggle switches to form a toggle triad. Our simulations show that this toggle triad can lead to co-existence of cells into three differentiated ‘single positive’ phenotypes – (high A, low B, low C), (low A, high B, low C), and (low A, low B, high C). Moreover, the hybrid or ‘double positive’ phenotypes – (high A, high B, low C), (low A, high B, high C) and (high A, low B, high C) – can co-exist together with ‘single positive’ phenotypes. Including self-activation loops on A, B and C can increase the frequency of ‘double positive’ states. Finally, we apply our results to understand the cellular decision-making in terms of differentiation of naïve CD4+ T cells into Th1, Th2 and Th17 states, where hybrid Th1/Th2 and hybrid Th1/Th17 cells have been reported in addition to the Th1, Th2 and Th17 ones. Our results offer novel insights into the design principles of a multistable network topology and provides a framework for synthetic biology to design tristable systems.

## Introduction

Elucidating the operating principles of complex regulatory networks driving cellular decision-making is a central question in dynamical systems biology. A central tenet involved in decision-making is the ability of cells to exhibit more than one stable states (phenotypes) in response to varying intracellular and/or extracellular conditions, without altering their genetic content. This feature is called as multistability (co-existence of more than one stable states/phenotypes) and is implicated in cellular differentiation and reprogramming [1]. Thus, decoding the emergent dynamics of multistable biological networks hold great promise not only for mapping the cellular differentiation paths, but also for synthetic biology and regenerative medicine applications [2,3].

A commonly observed network motif involved in enabling multistability is a toggle switch, i.e. a set of two opposing transcription factors A and B, each of which is a master regulator of a specific cell-fate and can inhibit the activity of the other through direct or indirect mechanisms. This mutual repression can allow for two states – (high A, low B) and (low A, high B), and drives the ‘either-or’ choice between two cell-fates for a common progenitor cell [2]. For instance, in hematopoietic stem cells, mutual repression between PU.1 and GATA1 can drive a common myeloid progenitor to a myeloid cell fate (high PU.1, low GATA1) or an erythroid one (low PU.1, high GATA1) [2]. This mutual exclusion of the two master regulators in the two states is critical for establishing stable cellular identities, and can shepherd an ‘all-or-none’ response instead of a graded one [4– 6]. The construction of a toggle switch synthetically in *E. coli* set the stage for synthetic biology, when cells were shown to exhibit two states (bistability) and the ability to flip between them in response to transient stimuli [7]. Toggle switches and bistability is present in diverse biological contexts [8–10], and have received enough theoretical attention for their dynamics too [11–14].

One or both of the two master regulators in a toggle switch (A and B) can self-activate. Such self-activating toggle switches can allow for the existence of one more stable state (medium A, medium B) in addition to the two driven by a toggle switch. This third state often corresponds to a common progenitor cell-state, as seen across many instances of cellular differentiation [2,15]. This ‘intermediate’ progenitor state is often ‘metastable’ and can differentiate to one of the two relatively more stable terminal states [2]. However, the dynamics of networks giving rise to three distinct states with a common progenitor have not been relatively as well-studied [16], despite instances of such decision-making seen in differentiation of CD4 expressing T-cells [17,18].

Here, we investigate the emergent dynamics of a set of three mutually repressing master regulators (A, B and C), and show that this ‘toggle triad’ can lead to the co-existence of three distinct differentiated or ‘single positive’ phenotypes – (high A, low B, low C), (low A, high B, low C), and (low A, low B, high C). In addition, the three ‘double positive’ states – (high A, high B, low C), (high A, low B, high C), and (low A, high B, high C) – can also be seen to co-exist with ‘single positive’ phenotypes, although at a lower frequency. Adding self-activation on these master regulators can enrich for the existence of these ‘double positive’ phenotypes that can be thought of as intermediate cell states between the corresponding ‘single positive’ or differentiated states. Our results offer a mechanistic explanation of how a ‘toggle triad’ formed among RORγT, GATA3 and T-bet can allow for three distinct T-cell states – Th1 (high T-bet, low GATA3, low RORγT), Th2 (low T-bet, high GATA3, low RORγT), Th17 (low T-bet, low GATA3, high RORγT) as well as corresponding hybrid cell fates originating from a common progenitor states (naïve CD4^+^ T cell).

## Results

### Toggle triad can allow for co-existence of three phenotypes (tristability)

The emergent dynamics of simple two-component and three-component networks such as toggle switch and repressilator has been well-investigated [11–14,19–21]. A toggle switch (i.e. a set of two mutually repressing transcription factors) (**Fig 1A**) can lead to two phenotypes – (high A, low and (low A, high B), thus A/B << 1 or A/B >> 1 for the two stable states (**Fig 1B**). This stark difference in the relative levels of A and B in the two states can drive cellular differentiation, as seen in multiple scenarios during embryonic development [2]. These two phenotypes may co-exist (bistable region) for certain range of parameters (green shaded region in **Fig 1C**); however, tuning the levels of various cell-intrinsic or cell-extrinsic signals can lead to one of the states being destabilized, thus leading to two different monostable regions (pink shaded regions in **Fig 1C**).

**Figure 1:**
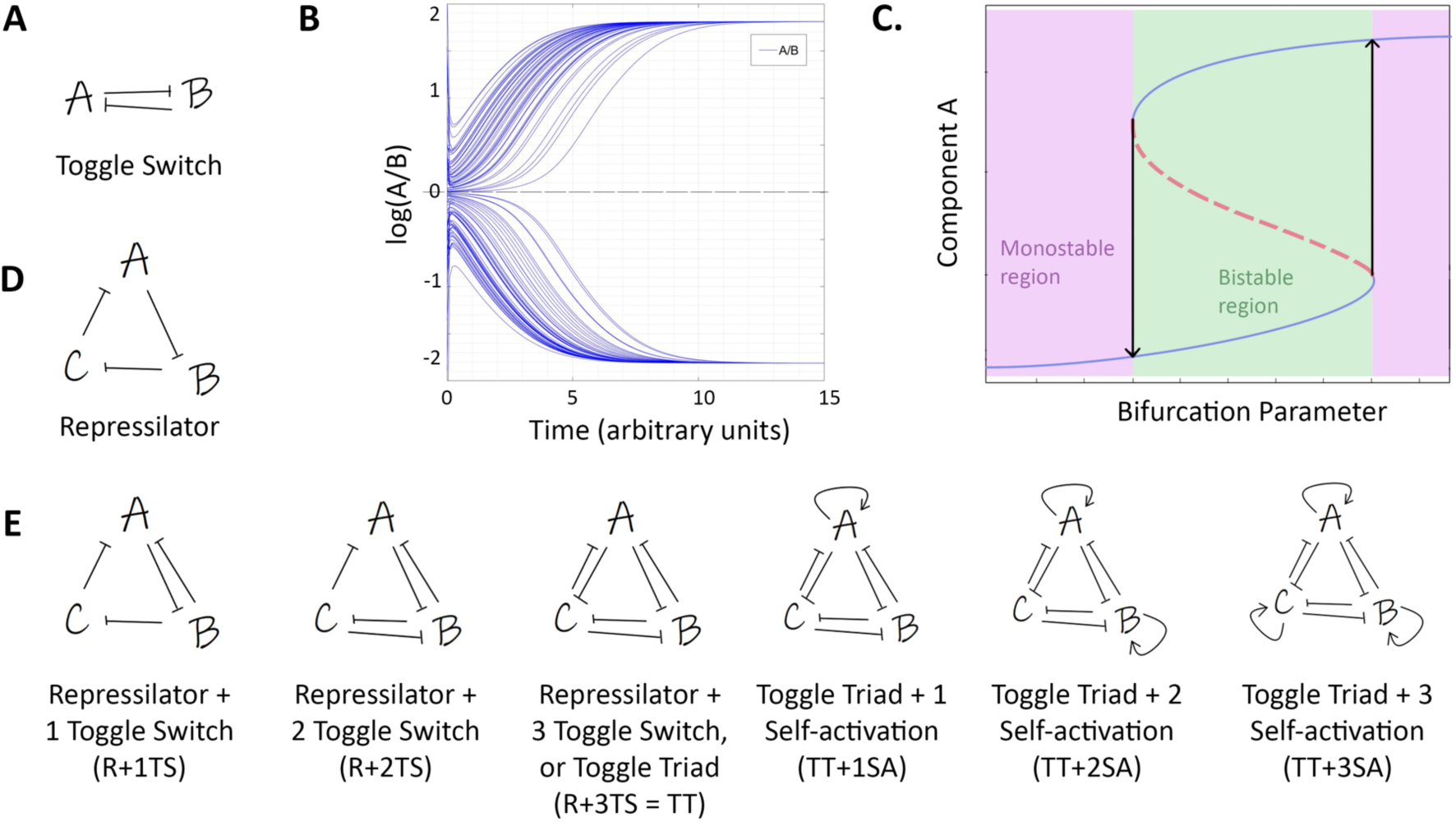
Network schematics. **A)** Toggle Switch. **B)** Dynamics of a toggle switch – different initial conditions can lead to two states: A/B >> 1 or A/B << 1. **C)** Schematic of the bifurcation diagram of a toggle switch. Solid blue curves indicate stable states; red dotted curves indicate unstable states. Bidirectional arrows show transition among different states. Green shaded region shows bistable region; pink shaded regions show two possible monostable regions. **D)** Schematic of a repressilator. **E)** Schematics of network topologies – repressilator with one, two, or three toggle switches (R+1TS, R+2TS, R+ 3TS = toggle triad), toggle triad with one, two, or three self-activations (TT+1SA, TT+2SA, TT+3SA).

A repressilator (**Fig 1D**; a cyclic arrangement of three inhibitory transcription factors), on the other hand, does not lead to multiple stable states, instead can display sustained or damped oscillations. Here, we investigate the dynamics of various possible couplings between the topology of these two well-characterized network motifs in different three-component systems. In a repressilator topology, either one (R+ 1TS), two (R+ 2TS) or three (R+3TS) toggle switches were integrated. The network formed by a set of three mutually repressing regulators is hereafter referred to as a ‘toggle triad’ (TT). Further, one, two or three of these regulators have been considered to be self-activatory as well (TT + 1SA; TT + 2 SA; TT+ 3 SA) (**Fig 1E**).

Next, to investigate the robust dynamical features of the abovementioned network topologies, we used a recently developed computational tool – RACIPE [22]. RACIPE takes the network topology as an input and converts it into a set of coupled ordinary differential equations (ODEs) to represent the set of interactions in that network topology. RACIPE samples 10,000 sets of parameters within a biologically relevant range, i.e. it generates an ensemble of mathematical models, each with a different parameter set. For every chosen parameter set, RACIPE chooses a random set of initial conditions for each node in the network, simulates the dynamics, and reports the different possible steady-state values for each node. Thus, each parameter set or kinetic model simulated via RACIPE corresponds to a different combination of parameters, reflecting cell-to-cell heterogeneity in biochemical reaction rates. An ensemble of models denotes the behavior of a cell population and statistical tools are then applied to identify the robust dynamic properties of the given network.

Here, each kinetic model is a set of three coupled ODEs, each of which tracks the dynamics of the levels of three interconnected molecular players A, B and C in various network topologies. Each of them have innate rates of production and degradation; the net production rate is affected by transcriptional regulation from other nodes; for instance, the inhibition of B by A in repressilator (R), repressilator + 1 toggle switch (R+1TS), repressilator + 2 toggle switches (R + 2 TS), and the toggle triad (TT) (**Fig 1D**). The set of differential equations is solved numerically to attain steady-state values of each node. For each given parameter set, depending on the initial condition, each of these molecular players can converge to one or more possible steady states enabled by the given parameter set. Thus, a circuit considered can be potentially multi-stable (i.e. two or more phenotypes).

For all seven network topologies – R, R + 1TS, R + 2 TS, TT, TT + 1 SA, TT + 2SA, TT + 3 SA (**Fig 1D**), we use RACIPE to quantify the number of parameter sets that led to only one phenotype (monostable) as well as those that led to two (bistable) and three (tristable) phenotypes. A repressilator has been shown to be capable of generating sustained or damped oscillations but not multistability; thus, as expected, the parameter sets generated by RACIPE enabled either monostability (damped oscillations) or sustained oscillations (**Fig 2A, S1**). As we include more inhibitory links in the network topology, moving from a repressilator to a toggle triad, the frequency of parameter sets leading to monostability decrease continuously, and those leading to multistable solutions – either bistable or tristable – increase (**Fig 2A**). Next, we investigate the dynamics of toggle triad with one or more self-activations included (TT, TT + 1SA, TT + 2SA, TT + 3SA) via RACIPE (**Fig 2B**). A toggle triad has ∼53% of parameter sets driving monostability; this frequency sharply decreases as one or more self-activations were included in the network topology. Instead, the frequency of parameter sets enabling tristability monotonically increase as we add more self-activations; while that of parametric combinations corresponding to bistability increases initially but decreases again (**Fig 2B**). Put together, a toggle triad – with or without one or more self-activation links (TT, TT + 1SA, TT + 2 SA, TT + 3SA) – is capable of exhibiting tristability, i.e. co-existence of three distinct stable states (phenotypes).

**Figure 2:**
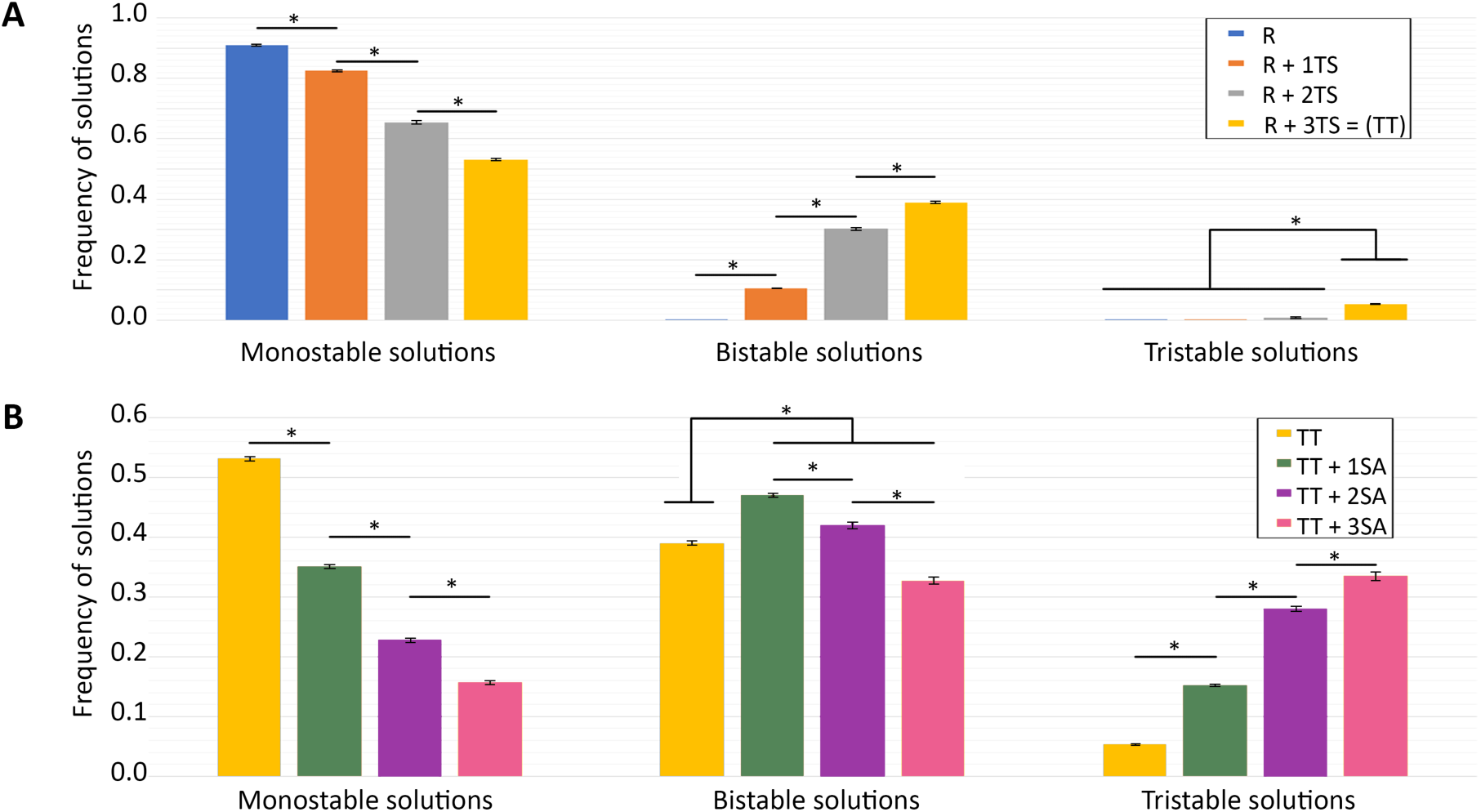
RACIPE outputs for networks shown in Fig 1E. **A, B)** Frequency of parameter sets used by RACIPE that enable monostable, bistable, tristable solutions for different networks. N=3 independent RACIPE replicates were done; error bars denote standard deviation. * denotes p <0.05 for a Students’ t-test.

### Toggle triad can enable three predominant states – (high A, low B, low C), (low A, high B, low C) and (low A, low B, high C)

We next characterized the different steady states/phenotypes that a toggle triad can allow for, as identified by RACIPE. A toggle triad allows for ∼53% monostable cases, ∼39% bistable cases, and ∼5% tristable cases (**Fig 3A**). We collated the levels of A, B and C obtained from all parameter combinations obtained via RACIPE and plotted them as a heatmap. This heatmap revealed three predominant states – (high A, low B, low C), (low A, high B, low C) and (low A, low B, high C) – represented by {Abc}, {aBc} and {abC} respectively hereafter. In addition to these states, a few instances of (high A, high B, low C), (high A, low B, high C) and (low A, high B, high C) – denoted by {ABc}, {AbC}, and {aBC} states respectively hereafter – were also observed (**Fig 3B**). These results indicate that a toggle triad can enable for states with one of the master regulators being relatively higher than the other two (‘single positive’ states), as well as those with two master regulators being relatively higher than the third one (‘double positive’ states).

**Figure 3:**
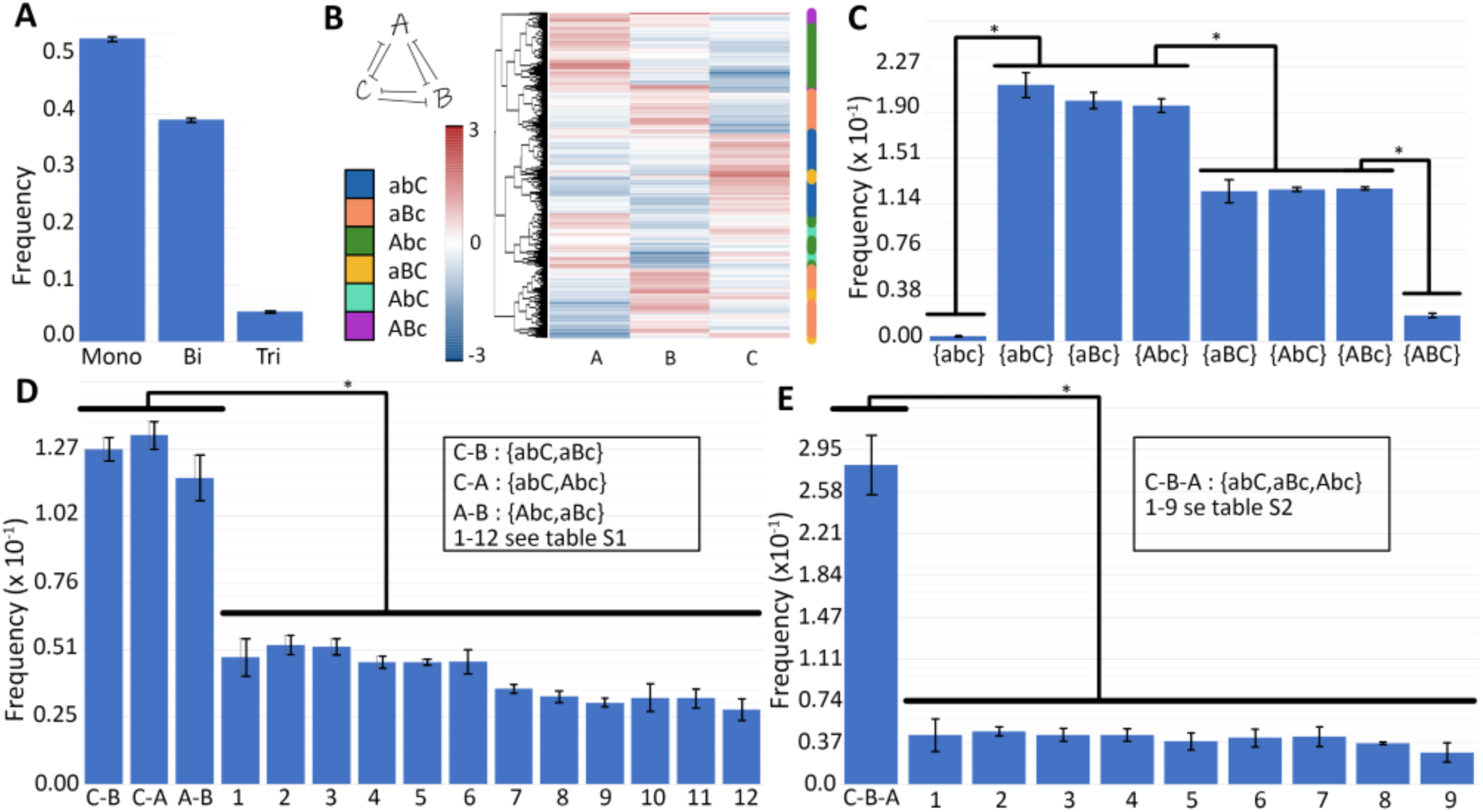
Characterization of a toggle triad. **A)** Frequency of monostable, bistable, tristable solutions for a toggle triad. **B)** Heatmap showing the monostable solutions for a toggle triad; the nomenclature shown capitalizes the node whose levels are relatively high. Thus, Abc denotes (A-high, B-low, C-low}, aBc denotes (A-low, B-high, C-low), abC denotes (A-low, B-low, C-high) (three ‘single positive’ states). Abc denotes (A-high, B-high, C-low), AbC denotes (A-high, b-low, C-high), and aBC denotes (A-low, B-high, C-high) (three ‘double positive’ states). ABC denotes (A-high, B-high, C-high) (triple positive), abc denotes (A-low, B-low, C-low) (triple negative) states. **C)** Frequency of 8 = (2^3) possible monostable solutions. **D, E)** Frequency of different bistable and tristable cases; with the most frequent ones being combinations of Abc, aBc and abC = {Abc, aBc}, {aBc, abC}, {abC, Abc} (bistable) and {aBc, Abc, abC} (tristable). Error bars represent the standard deviation over n=3 independent replicates of RACIPE. *: p<0.05 for Students’ t-test.

Reinforcing the trends seen in the heatmap, we observed that the distributions of levels of each of the three players A, B and C obtained via all RACIPE solutions was largely bimodal (**Fig S2**), indicating that each node in the network can exist in either a “high” or a “low” state. The minima of the distribution was around zero, thus it was chosen as the cutoff for defining “high” vs. “low” expression (see Methods). Together, a total of 8 (= 2^3^) states can exist for a toggle triad. Further, we quantified the relative frequency of these 8 possible steady states among monostable solutions. Among the parameter sets enabling monostable solutions, ∼60% cases led to the ‘single positive’ {Abc}, {aBc}, {abC} states, ∼36% of cases led to the ‘double positive’ {ABc}, {aBC}, {AbC} states, while only 4% of the cases led to ‘triple positive’ (high A, high B, high C - {ABC}) or ‘triple negative’ (low A, low B, low C – {abc}) states (**Fig 3C**). Owing to the symmetric nature of the toggle triad, there was striking symmetry in the number of parameter cases leading to each of the three ‘single positive’ or ‘double positive’ states as well; i.e. ∼60/3 = 20% of parameter sets each converged to {Abc}, {aBc} or {abC} as a steady state, and ∼36/3 =12% of parameter sets converged to {ABc}, {aBC} or {AbC} as a steady state (**Fig 3C**). Given the negligible frequency of the ‘triple positive’ and the ‘triple negative’ solutions, they were excluded from our further analysis.

We next examined the frequency distribution of parameter sets leading to bistable solutions. Here, a total of 15 (=^6^C_2_; number of ways to choose two out of six solutions) phases (i.e. combinations of steady states). The three most common phases were the ones with co-existing ‘single positive’ states, i.e. {Abc, aBc}, {aBc, abC} and {Abc, abC}, totaling up to ∼42% of all parameter sets (**Fig 3D, Table S1**). The remaining 12 combinations of steady states were obtained from a cumulative ∼56% of parameter sets enabling bistability. Similarly, among a total of 20 (=^6^C_3_; number of ways to choose three out of six solutions) tristable solutions, the most frequent combination was the set of co-existing ‘single positive’ states, i.e. {Abc, aBc, abC} (∼30% parameter sets) (**Fig 3E, Table S2**). Put together, these results suggests that one of the underlying design principles of a toggle triad is to allow the existence (or co-existence) of phenotypes where the levels of one of the three components is much larger than the remaining two, i.e. both A/B>> 1 and A/C>>1 (i.e. {Abc}), or both B/A>>1 and B/C>> 1 (i.e. {aBc}) or both C/A>> 1 and C/B>>1 (i.e. {abC}).

### The dynamical traits of a toggle triad

To further test that the abovementioned design principles are specific to the toggle triad topology, we investigated the dynamics of multiple three-component networks where one or more of the six inhibitory links in a toggle triad has/have been replaced with an activatory link (circuits C1-C14; **Fig S3**). All of the 14 circuits failed to exhibit at least one of the salient features of toggle triad, when comparing the state frequency for parameter sets enabling monostable solutions: a) the frequency of ‘triple positive’ and ‘triple negative’ states is negligible (8/14 cases have 18% or more parameter sets leading to either of these two states), b) the relative frequency of all three ‘single positive’ states was the same, c) the relative frequency of all three ‘double positive’ states was the same, and d) the ‘single positive’ states were more frequent than the ‘double positive’ ones (**Table S3**). As expected, the circuit with all inhibitory links in a toggle triad replaced by activation (C2) showed the ‘triple negative’ and ‘triple positive’ states as the most predominant ones. Similarly, when comparing the results for bistable and tristable scenarios, none of the 14 circuits showed the co-existence of 2 (in case of bistable) or 3 (in case of tristable) ‘single positive’ cases as the predominant trend as seen in case of a toggle triad (**Tables S4, S5**). Finally, the percentage distribution of parameter sets that led to monostable, bistable and tristable solutions were quite different for these 14 circuits as compared to a toggle triad (**Table S6, Fig S4-S9**). To gain further confidence in these results via RACIPE, we simulated the dynamics of toggle triad and circuit C2 using asynchronous Boolean modeling approach [23] – a parameter-independent approach – and observed similar trends as seen in RACIPE, suggesting the key role of network topology instead of specific parametric combinations in enabling these robust design dynamical principles of a toggle triad (**Table S7**). Overall, these results strengthen the association of a toggle triad formed by A, B and C with the existence/co-existence of these states – (high A, low B, low C), (low A, high B, low C) and (low A, low B, high C).

To further characterize the parameter space for the co-existence of states, we drew bifurcation diagrams for multiple parameter sets identified via RACIPE that enabled the three most frequent bistable phases - {Abc, aBc}, {aBc, abC} and {Abc, abC}. For a representative parameter set pertaining to the bistable phase enabling (low A, high B, low C) and (low A, low B, high C) (i.e. {aBc, abC}), the degradation rate of B (kb) was chosen as bifurcation parameters. Over a wide range of parameter values of kb spanning an order of magnitude (0.01 – 0.4), both the states aBc and abC were observed to co-exist. At very high values of kb, the (low A, high B, low C) state, i.e. aBc was no longer seen; instead, the system had only the (low A, low B, high C) state (**Fig 4A-C**). It was encouraging to see that this trend of bistability was qualitatively maintained even when the degradation rate of C (i.e. kC) was varied by +/- 25% relative to the one identified by RACIPE (**Fig 4A,B**), thus suggesting that the co-existence of more than one such ‘single positive’ state can be a robust dynamical feature of a toggle triad.

**Figure 4:**
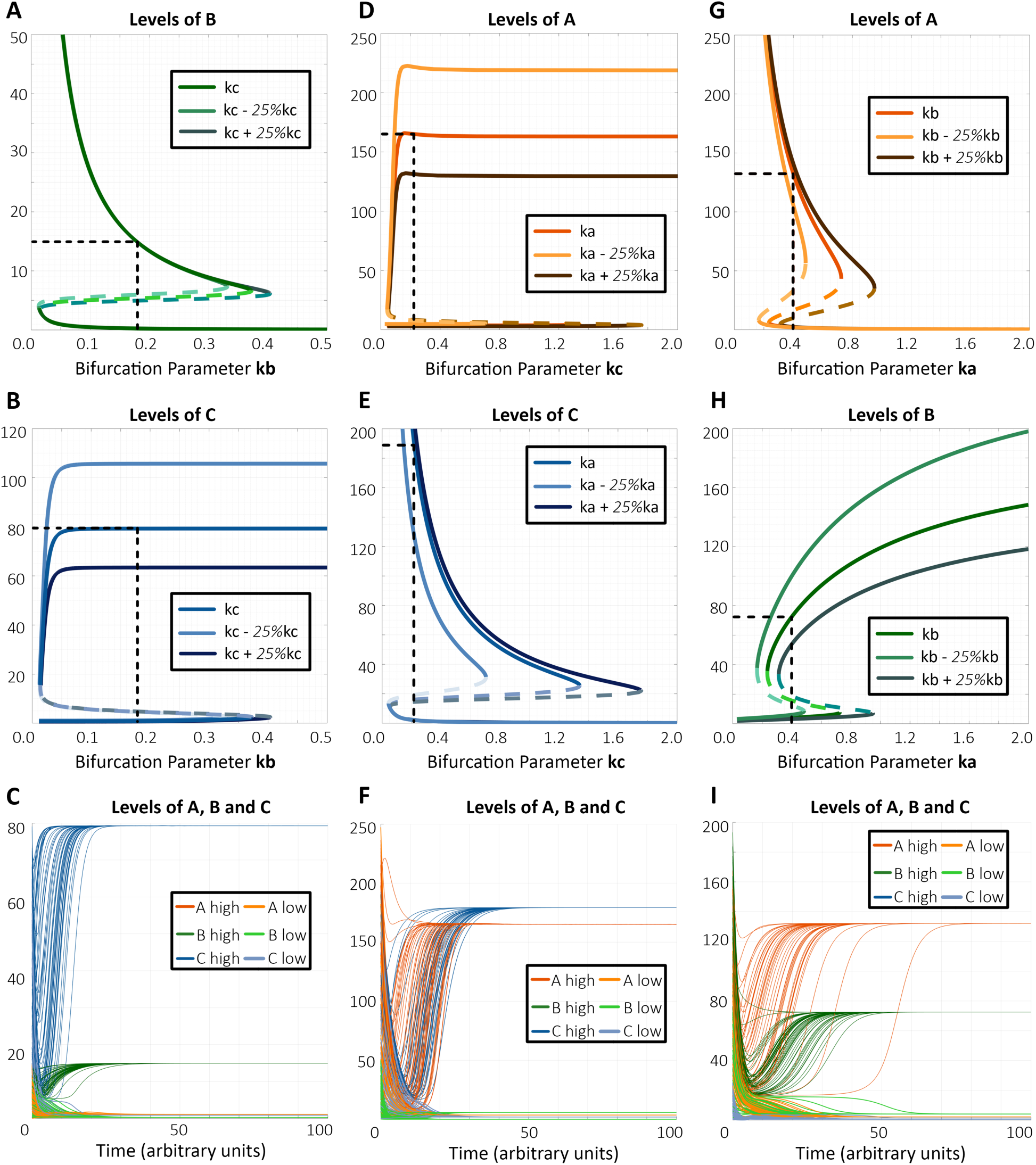
Bifurcation diagrams and dynamics plots for representative cases of bistable phases. **A)** Bifurcation diagram of expression level of component B with kb as bifurcation parameter for the bistable phase {aBc, abC}. **B)** Same as A) but for component C. **C)** Dynamics plots of expression levels of components A, B and C for the bistable phase {aBc, abC}, showing convergence to two different states with varied levels of B and C (levels of A are low in both cases). **D)** Bifurcation diagram of expression level of component A with kc as bifurcation parameter for the bistable phase {Abc, abC}. **E)** Same as D but for component C. **F)** Dynamics plots of expression levels of components A, B and C for bistable phase {Abc, abC}, showing convergence to two different states with varied levels of A and C (levels of B are low in both cases). **G)** Bifurcation diagram of expression level of component A with ka as bifurcation parameter for the bistable phase {aBc, Abc}. **H)** Same as G) but for component B. **I)** Dynamics plots of expression levels of components A, B and C for the bistable phase {aBc, Abc}, showing convergence to two different states with varied levels of A and B (levels of C are low in both cases). Parameter values for columns A-C, D-F and G- I are given correspondingly in Table S13. The bifurcation diagrams for ‘low’ components (component A in panels A-C, component B in panels D-F, component C in panels G-I) are shown in Fig S11.

Similarly, for the bistable phase enabling (high A, low B, low C) and (low A, low B, high C) states (i.e. {Abc, abC}), these states co-existed over a wide range of parameter values of kc spanning an order of magnitude (0.05 – 1.8) (**Fig 4D-F**). This robust behavior was maintained for scenarios when the degradation rate of A (i.e. kA) was either increased or decreased by 25% of the value identified via RACIPE (**Fig 4D, E**). Consistent results were observed for the {Abc, aBc} state as well (**Fig 4G-H**), as well as additional parameter sets for these three most common bistable phases (**Fig S10, S11**). Put together, these results reveal that the parameter space associated with a toggle triad can allow for co-existence of ‘single positive’ states and possible switching among them under the influence of intrinsic noise.

Next, to better decode the co-existence of ‘single positive’ and ‘double positive’ states, we performed similar bifurcation analysis for the bistable phases containing one ‘single positive’ and one ‘double positive’ state. For a representative parameter set pertaining to the bistable phase enabling {aBC, abC}, we performed bifurcation analysis for the degradation rate of C (i.e. kc), and observed bistability over an order of magnitude (kc= 0.1-1.4; **Fig 5A, B**). The bistable nature was conserved for +/-25% variations in kb. Similar patterns were seen for {Abc, AbC} (**Fig 5C,D**) and {aBc, ABc} states (**Fig 5E,F**).

**Figure 5:**
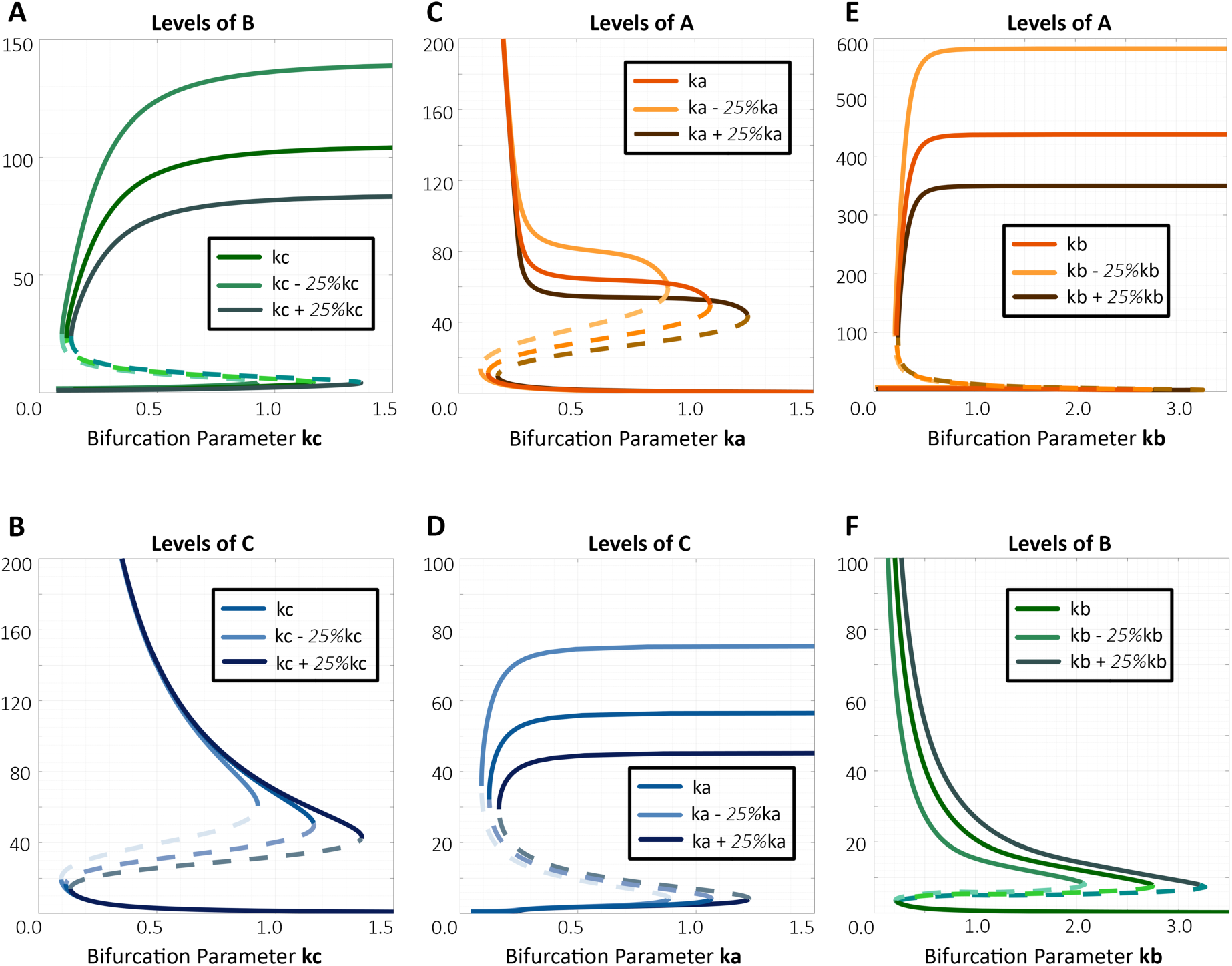
Bifurcation diagrams for bistable phase with combination of a ‘single positive’ state and a ‘double positive’ state. **A)** Bifurcation diagram of expression level of component B with kc as bifurcation parameter for the bistable phase {aBC, abC}. **B)** Same as A) but for component C. **C)** Bifurcation diagram of expression level of component A with ka as bifurcation parameter for the bistable phase {Abc, AbC}. **D)** Same as C) but for component C. **E)** Bifurcation diagram of expression level of component A with kb as bifurcation parameter for the bistable phase {aBc, ABc}. **F)** Same as E) but for component B. Corresponding parameter sets for columns A-B, C-D and E-F are given in Table S13.

Further, we chose a representative parameter case corresponding to tristable phase containing ‘single positive’ states – {Abc, aBc, abC}. For parameters identified via RACIPE, the system converged to three distinct steady states (**Fig S12A**). Altering the values of kb and/or kc disrupted tristability and led to the three monostable regions – (low A, high B, low C) (low kb; **Fig S12B**), (low A, low B, high C) (low kc; **Fig S12C**), and (high A, low B, low C) (high kb and kc; **Fig S12D**). Similar results were seen for another parameter set obtained via RACIPE for tristability (**Fig S13**). These results suggest that the tristability of a toggle triad – co-existence of three ‘single positive’ states – can be disrupted if one of the three components has a very different stability (i.e. half-life) as compared to others.

### Toggle triad with self-activations enrich for the existence of ‘double positive’ states

Next, we probed the dynamics of a toggle triad with self-activations on all three nodes (A, B, C) (TT + 3SA). We collated the steady-state levels of A, B and C obtained from all parameter combinations obtained via RACIPE for this circuit and plotted them as a heatmap. Similar to the case of a toggle triad, the ‘single positive’ (high A, low B, low C), (low A, high B, low C) and (low A, low B, high C) states were predominant. However, as compared to a toggle triad, there was a marked enrichment of the ‘double positive’ states - (high A, high B, low C), (high A, low B, high C) and (low A, high B, high C), i.e. {ABc}, {AbC}, and {aBC} states respectively (**Fig 6A**). Furthermore, in the case of a toggle triad with self-activations, the number of parameter sets enabling monostability was 3.3 fold lower as compared to a toggle triad (53% for TT vs. 16% for TT + 3 SA) and a 6.8 fold increase in those enabling tristability (5% for TT vs. 34% for TT + 3SA) (**Fig 6B-C**). The toggle triad with self-activation also exhibited tetrastable, pentastable and hexastable behavior (13% parameter sets for tetra-stability, 4% for penta-stability) (**Fig 6C**).

**Figure 6:**
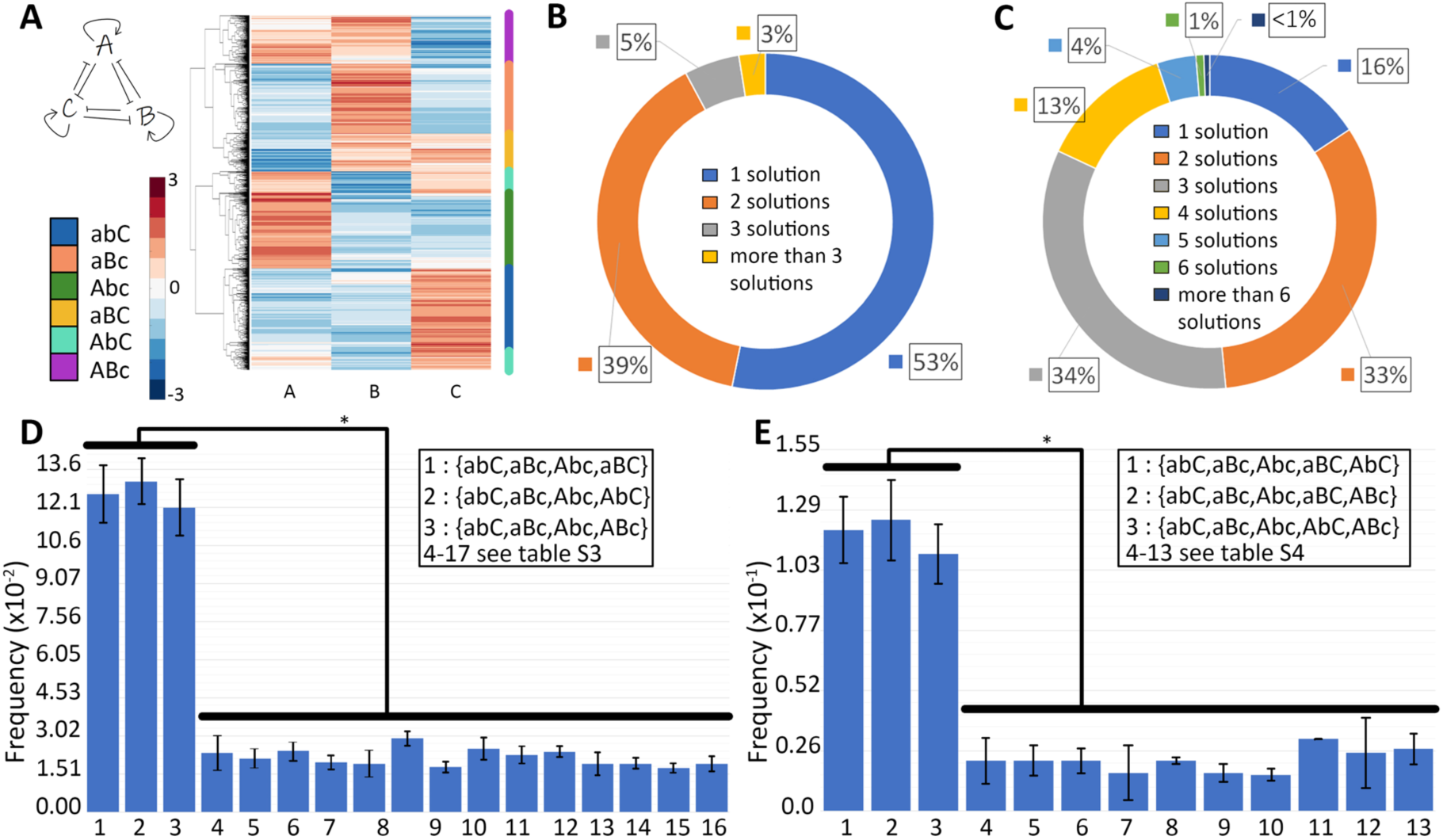
Characterization of toggle triad with three self-activations (TT + 3SA). **A)** Heatmap showing monostable solutions for TT+3SA; Heatmap showing the monostable solutions for a toggle triad; the nomenclature shown capitalizes the node whose levels are relatively high. Thus, Abc denotes (A-high, B-low, C-low}, aBc denotes (A-low, B-high, C-low), abC denotes (A-low, B-low, C-high) (three ‘single positive’ states). Abc denotes (A-high, B-high, C-low), AbC denotes (A-high, b-low, C-high), and aBC denotes (A-low, B-high, C-high) (three ‘double positive’ states). **B)** Frequency of monostable, bistable, tristable solutions in a toggle triad (TT) shown as a pie chart. **C)** Frequency of monostable, bistable, tristable, tetrastable, pentastable solutions for a TT+3SA case shown as a pie chart. **D, E)** Frequency of different tetrastable and pentastable phases are combinations of Abc, aBc, abC, aBC, AbC, ABc = {Abc, aBc, abC, aBC}, {Abc, aBc, abC, AbC}, {Abc, aBc, abC, ABc} (tetrastable) and {aBc, Abc, abC, aBC, AbC}, {aBc, Abc, abC, ABc, AbC}, {aBc, Abc, abC, aBC, ABc} (pentastable). Error bars denote the standard deviation of n=3 independent RACIPE simulations. * denotes statistical significance (p<0.05 for Students’ t-test).

The predominance of the ‘single positive’ states and their combinations prevailed in monostable, bistable and tristable parameter sets for the case of a toggle triad with three self-activations (**Table S8-10**). In the tetrastable cases, the top three most predominant combinations contained all the three ‘single positive’ states with one of the three possible ‘double positive’ states – {Abc, aBc, abC, ABc}, {Abc, aBc, abC, AbC} and {Abc, aBc, abC, aBC} (**Fig 6D; Table S11**). Similarly, in pentastable cases, the top three most predominant combinations contained all the three ‘single positive’ states with two of the three possible ‘double positive’ states – {Abc, aBc, abC, ABc, aBC}, {Abc, aBc, abC, AbC, aBC} and {Abc, aBc, abC, ABc, AbC} (**Fig 6E; Table S12**), unraveling the dynamical traits of a toggle triad where each of the master regulator can self-activate. Put together, these results suggest that a toggle triad with self-activation can enrich for the (co-) existence of such ‘double positive’ states.

So far, our analysis has focused on deterministic dynamics. Given that molecular fluctuations can have a profound impact on phenotypes chosen by the system [24], including those for a toggle switch [12], we analyzed the toggle triad with self-activation using sRACIPE (stochastic version of RACIPE) [25]. Stochastic simulations using sRACIPE shows spontaneous switching among different possible states – at least among all ‘single positive’ ones, across multiple parameter sets. ‘Double positive’ states are also seen albeit with lower residence times (**Fig 7A-C; Fig S14, S15**).

**Figure 7:**
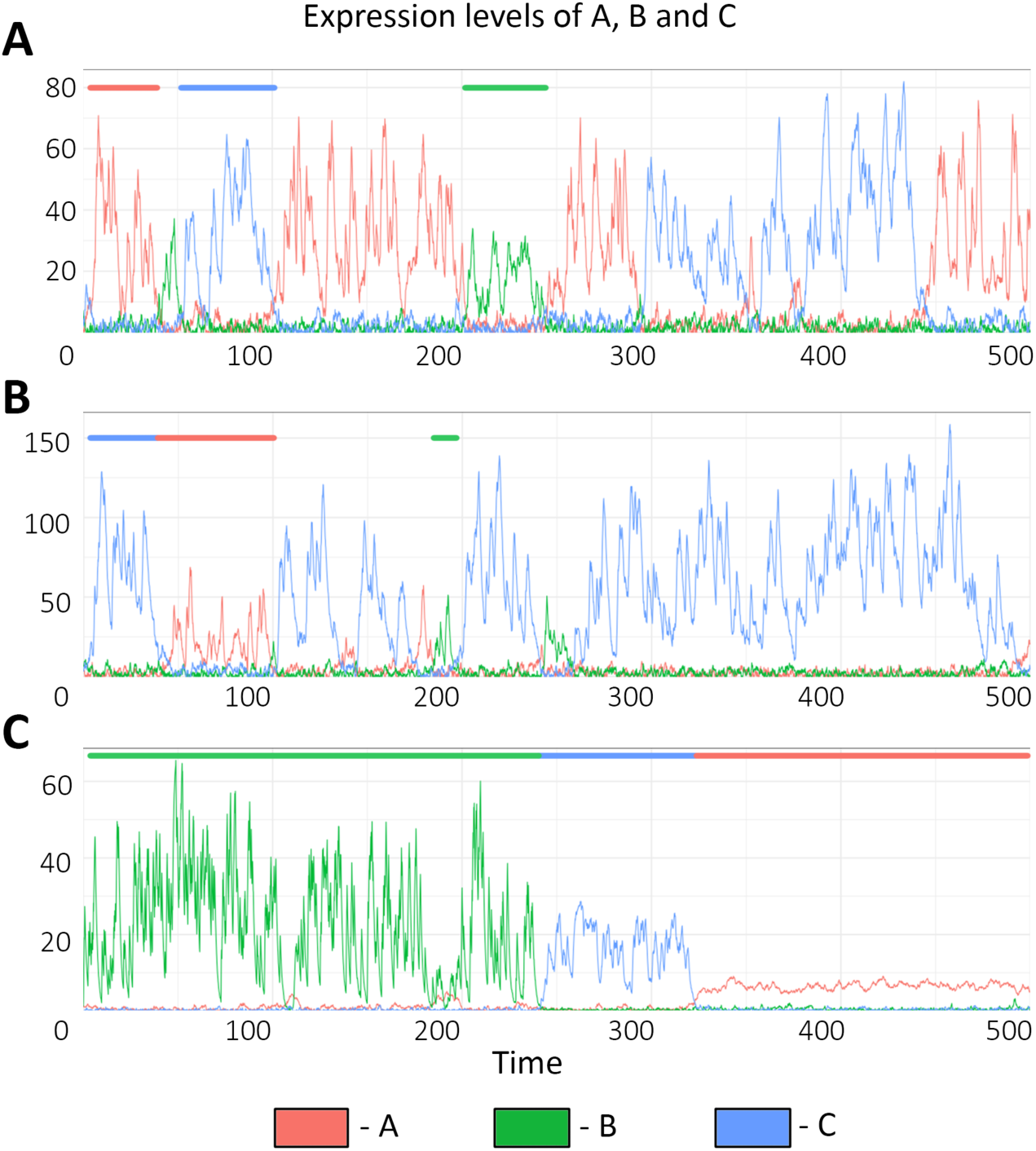
sRACIPE results for one of the replicates of three different parameter sets used. **A)** Dynamics plot showing switching between states for parameter set 1. Color bars on top representatively mark the regions of each of the states – green bar shows (low A, high B, low C), red bar shows (high A, low B, low C) and blue bar shows (low A, low B, high C). **B, C)** Same as A) but for a replicate of parameter set 2 and 3 respectively. The plots for two other replicates for each of the parameter sets are shown in Fig S14, S15. Parameter values for sRACIPE for all the three replicates and parameter sets are given in Table S13.

### Design principles of multistability enabled by toggle triad with/without self-activation

As discussed above, a toggle triad with/without self-activation can be monostable or multistable. Thus, we investigated different parameter combinations identified by RACIPE enabling the three most frequent monostable ({Abc}, {aBc}, {abC}), three most frequent bistable ({Abc, aBc}, {Abc, abC}, {aBc, abC}) and the most frequent tristable ({Abc, aBc, abC}) phases. We hypothesized that the relative strengths of different regulatory links in the network led to these different phases. In RACIPE formulation, the effect of inhibition from one node to another is captured by a shifted Hill function described by three independent parameters: n (cooperativity), λ (fold change), and H_0_/(g/k) (relative half-maximal concentration or threshold) [22]. The higher the value of n, the stronger the repression, and the higher the value of H_0_/(g/k), the weaker the repression. For inhibitory links, λ varies between 0 (very strong repression) to 1 (no effect). Thus, the higher the value of x = n/(λ*H_0_/(g/k)), the stronger the corresponding inhibition. For parameter sets enabling {Abc}, i.e. (high A, low B, low C), we hypothesized that the inhibition of B and C by A is stronger than the inhibition of A by B and C. To test this hypothesis, we quantified the values of x for A inhibiting B (x (A –|B)) and those of B inhibiting A (x (B –|A)) for all parameter sets enabling {Abc}. We found that for 90% of parameter sets, x (A –|B) > x (B –|A) (**Fig 8A**, columns 1 and 2; **Fig 8B**); similar trend was seen for toggle switch between A and C, i.e. x (A –|C) > x (C –|A) (**Fig 8A**, columns 3 and 4). However, no such trend was seen for toggle switch between B and C (**Fig 8A**, columns 5 and 6). The large degree of overlap seen in distributions for x (A –|B)/x (B –|A) and for x (A –|C)/x (C –|A) suggest that in most parameter sets enabling (high A, low B, low C) state, B and C are both simultaneously strongly inhibited by A (**Fig 8B**). Consistent corresponding trends were seen for parameter sets enabling {aBc} and {abC} monostable phases (**Fig S16A, B, D, E**).

**Figure 8:**
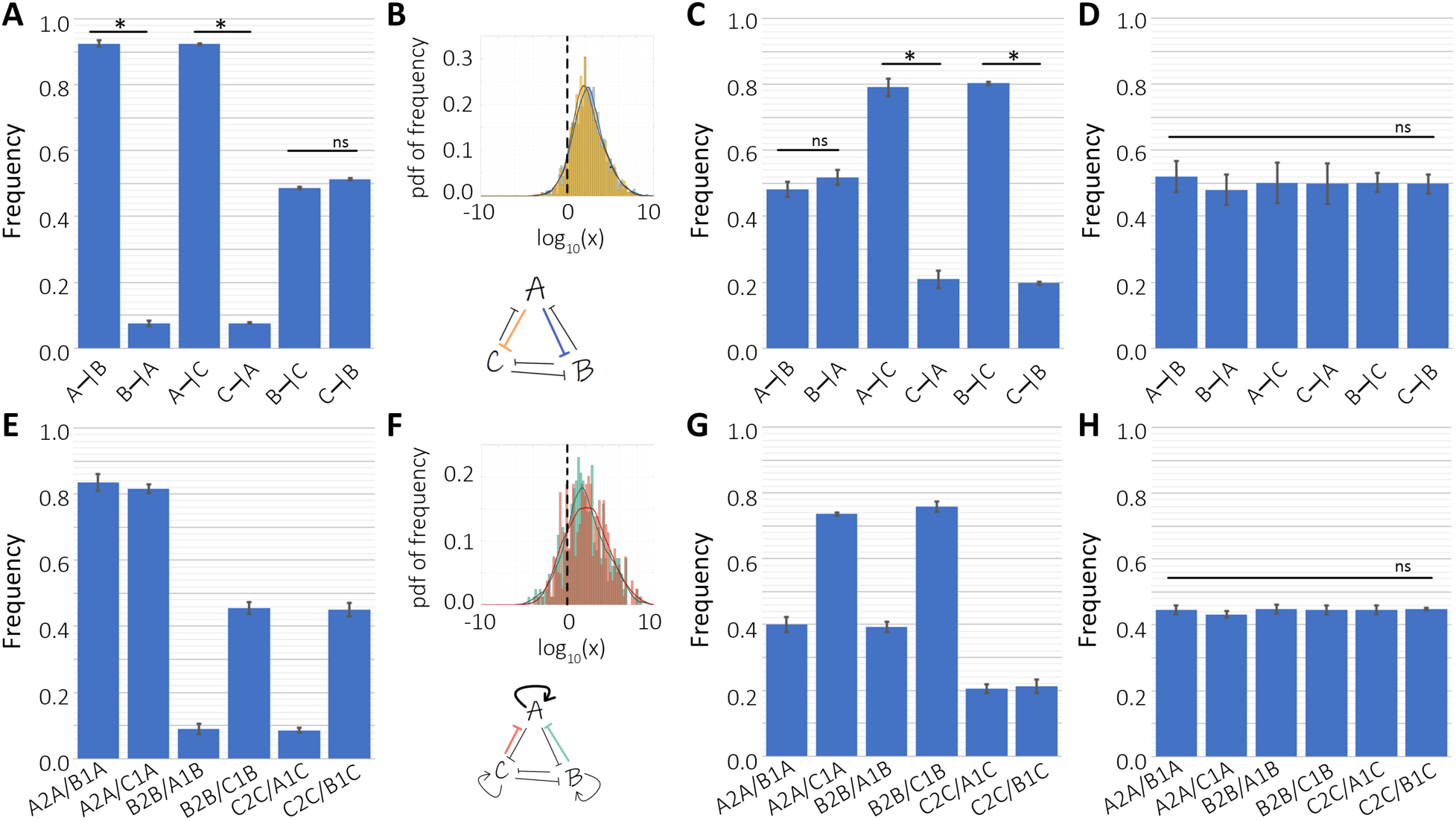
Design principles of toggle triad (TT) and that with self-activation (TT+3SA). **A)** Frequency of dominance of six inhibitory links for the case of monostable (A high, B low, C low) (i.e. {Abc}) in TT. X –|Y denotes the frequency of parameter sets when the inhibition of X on Y was stronger than the that of Y on X; thus adjacent two columns total up to 1. **B)** (top) Probability distribution functions of histograms of frequency of values of x(A–|B)/x(B–|A) (blue) and values of x(A–|C)/x(C–|A) (yellow) in TT for monostable case {Abc}. The x-axis is log_10_ transformed and the dotted line represents the numerical value 1. (below) Schematic showing that A inhibiting B and C are stronger links than B and C inhibiting A overall for parameter sets corresponding to monostable {A} **C)** Same as A) but for bistable (i.e. {Abc, aBc}) in TT. **D**) Same as A) but for tristable case {Abc, aBc, abC}) in TT. **E)** Frequency of parameter cases for which self-activation dominates upon incoming inhibitory link, for monostable case {Abc} in TT+3SA. X2X/X1Y denotes the percentage of parameter sets for which X self-activation is stronger than Y inhibiting X. **F)** (top) Probability distribution functions of histogram of frequency of x(A2A)/x(B1A) and x(A2A)/x(C1A) in TT+3SA. The x-axis is log_10_ transformed and the dotted line represents the numerical value of 1. (bottom) Schematic showing that for most of the parameter sets corresponding to {Abc}, self-activation of A dominates inhibition of A by B or C. **G)** Same as E) but for bistable case {Abc, aBc} in TT+3SA. **H)** Same as E) but for tristable case {Abc, aBc, abC} in TT+3SA. * denotes statistical significance (p<0.01 for students’ t-test). ‘ns’ denotes statistically non-significant cases. Parameters corresponding to Fig8, S16 are given in Tables S14, S15.

For the bistable phase {Abc, aBc}, we hypothesized that inhibition of C by A and B is stronger than inhibition of A and B by C. Indeed, we saw that 80% of parameter sets had x (A –|C) > x (C –|A) and x (B –|C) > x (C –|B) (**Fig 8C**, columns 3-6) but the relative strengths of mutual inhibition between A and B were not skewed (**Fig 8C**, columns 1-2). Similar corresponding trends were seen for other bistable phases {Abc, abC}, {aBc, abC} (**Fig S16 C,F**). For the tristable phase {Abc, aBc, abC}, the distributions of relative strengths of inhibition in any of the three toggle switches was not imbalanced or skewed (**Fig 8D**) as seen for monostable or bistable cases, suggesting that tristability is enabled only when the strengths of mutual inhibition among all three components are relatively well-balanced.

In case of toggle triad with self-activation, we defined x = n*λ/(H_0_/(g/k)) for the self-activatory links because the corresponding values of λ are higher than 1, and the higher the value of λ, the stronger the self-activation. For parameter sets in this case enabling {Abc}, i.e. (high A, low B, low C), we hypothesized that the self-activation of A is stronger than inhibition of A by B and C. Indeed, for over 80% of parameter sets, x(A2A)/x(B1A) > 1 and x(A2A)/x(C1A) > 1, where x(A2A) represent the strength of self-activation, and x(B1A) and x(C1A) denote the strength of inhibition of A by B and C respectively (**Fig 8E**, columns 1,2). Reminiscent of trends seen for toggle triad, the values x(A2A)/x(B1A) and x(A2A/C1A) span multiple orders of magnitude, and their overlap suggests that the self-activation of A is stronger than both the inhibition by B and that by C incumbent on A (**Fig 8F**). Conversely, the number of parameter sets showing a stronger self-activation of B and C relative to their inhibition by A is less than 10% (**Fig 8E**, columns 3,5). The relative strengths of links in the self-activating toggle switch between B and C show no such imbalance in distribution as expected; 50% of parameters have x(B2B)/x(C1B) >1 and remaining 50% have x(B2B)/x(C1B) < 1 (**Fig 8E**, columns 4, 6).

For ∼75% of parameter sets enabling the bistable phase {Abc, aBc}, self-activations of A and B was found to be stronger than inhibition of A and B by C (**Fig 8G**, columns 2, 4). Conversely, the self-activation of C was weaker than inhibition of C by A and B (**Fig 8G**, columns 5, 6). As expected, the self-activatory toggle switch between A and B was found to be well-balanced (**Fig 8G**, columns 1, 3), thus enabling the co-existence of (high A, low B, low C) and (low A, high B, low C) states. Consistent corresponding trends were seen for other parameter sets and phases (**Fig S16G-L**).

For the parameter sets enabling a tristable phase {Abc, aBc, abC}, all three nodes displayed a delicate balance in relative strengths of inhibition by other two nodes, and self-activation (**Fig 8H**). Interestingly, the frequency of parameter sets for which self-activation was stronger than inhibition received was slightly smaller than 50% (**Fig 8H**), suggesting that strong self-activation may not enable tristability of these ‘single positive’ states. These results are reminiscent of observations for a self-activating toggle switch between A and B, where a very strong self-activation of A and B relative to mutual inhibition led to only the ‘double positive’ (or hybrid A/B) state, and the two ‘single positive’ states (high A, low B) and (low A, high B) disappeared [15]. Thus, strong self-activation on all three components may enable the co-existence of three ‘double positive’ states.

Put together, this analysis reveals the patterns in multi-dimensional parameter space which enable a toggle triad (with/without self-activation) to display such a wide varsity in dynamical behavior – existence of three distinct states and the co-existence of any two or all three states.

### Th1/ Th2/ Th17 cell differentiation: a case study of toggle triad

Upon activation, naïve helper T cells differentiate towards a specific helper T cell subset. In the presence of specific activating signals a majority of these cells differentiate towards a particular subset expressing a lineage-specific transcription factor (master regulator). However, a small but significant number of cells may also expressing multiple master regulators [26]. To understand if the presence of cells expressing multiple master regulators may be explained through the toggle triad system described above, we undertook the case study involving three helper T cell subsets - Th1 (T-bet), Th2 (GATA3) and Th17 (RORγT). Assuming that these master regulators may mutually repress each other [18] (**Fig 9A**), we projected the steady state solutions obtained from the heatmap for a toggle triad (**Fig 3A**) on the two-dimensional scatter plots for (T-bet, GATA3) axes, (T-bet, RORγT) axes and (GATA3, RORγT) axes. The plots showed the emergence of three clusters, each corresponding to a ‘single positive’ state – blue (high T-bet, low GATA3, low RORγT state), orange (low T-bet, high GATA3, low RORγT state) and green (low T-bet, low GATA3, high RORγT state) (**Fig 9B, S17, S18**), corresponding to Th1, Th2 and Th17 cell-fates, respectively. The hybrid ‘double positive’ (black dots) states were also observed in addition to the ‘single positive’ states – (high T-bet, high GATA3, high RORγT), (high T-bet, low GATA3, low RORγT) and (low T-bet, high GATA3, high RORγT), although at a lower frequency than the ‘single positive’ ones. These states can be mapped to hybrid Th1/Th2, Th2/Th17 and Th1/Th17 cell types.

**Figure 9:**
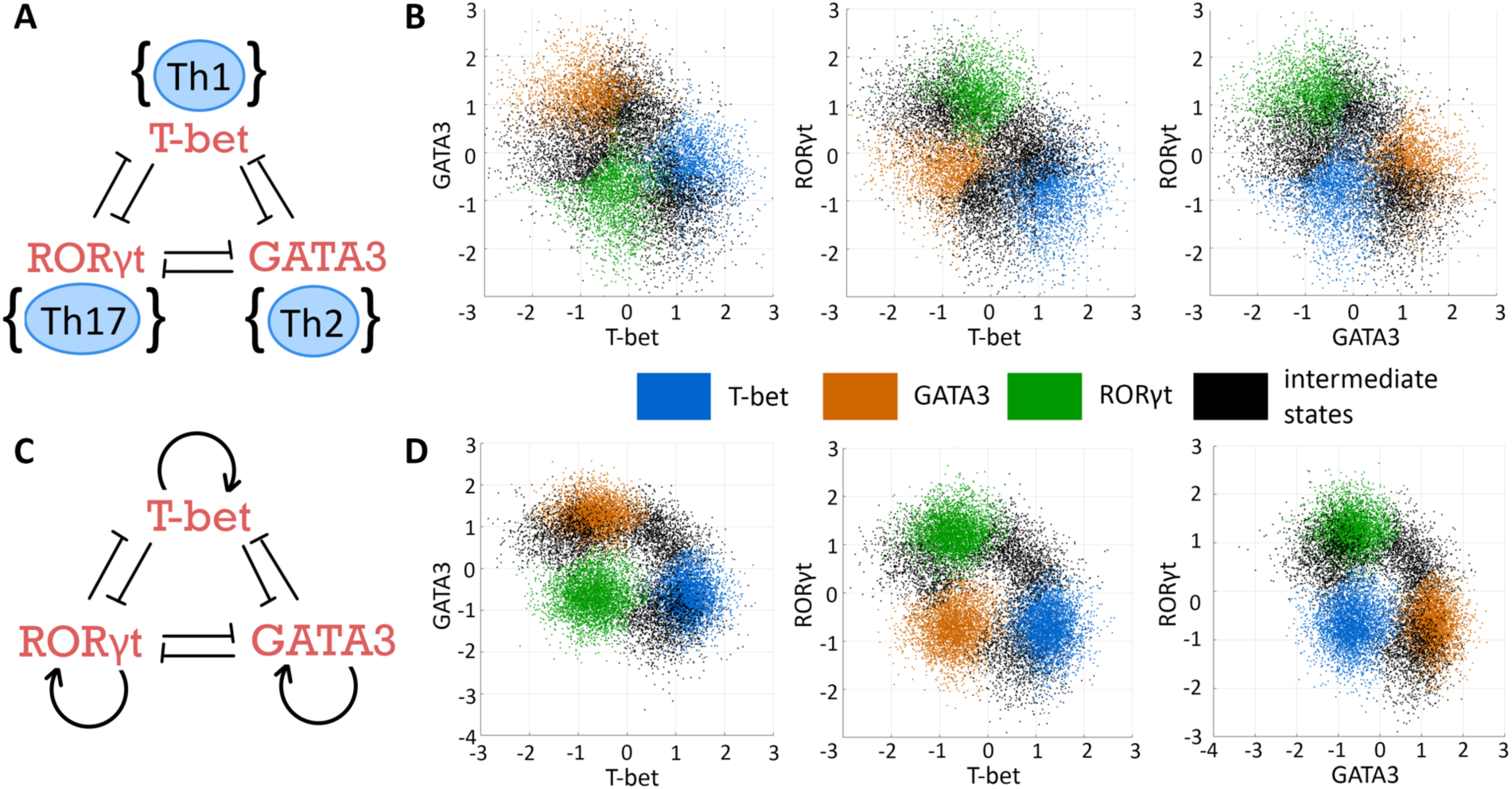
CD4 T-cell differentiation. **A)** Network showing proposed interaction among the master regulators of Th1, Th2 and Th17 – T-bet, GATA3 and RORγT – respectively. **B)** Two-dimensional scatter plots projecting solutions from the heatmap for a toggle triad network (Fig 3). **C)** Network of T-bet, GATA3 and RORγT including self-activations. **D)** Same as B but for solutions from the heatmap for a toggle triad with 3 self-activation (Fig 5). Blue colored dots denote Th1 (high T-bet, low GATA3, low RORγT), orange colored dots denote Th2 (low T-bet, high GATA3, low RORγT), green colored dots denote Th17 (low T-bet, low GATA3, high RORγT) state. Black dots denote the different hybrid states – Th1/Th2, Th2/Th17 and Th1/Th17: (high T-bet, high GATA3, high RORγT), (high T-bet, low GATA3, low RORγT) and (low T-bet, high GATA3, high RORγT) respectively. Data from respective heatmaps was subjected to k-means clustering to identify these six states (three ‘single positive’ and three ‘double positive’ ones).

GATA3, T-bet and RORγT have been found to self-activate directly and/or indirectly [27,28]. Thus, next, we included self-activation loops (**Fig 9C**), and projected the steady state solutions obtained from the heatmap for a toggle triad with self-activation (**Fig 5**) on the two-dimensional scatter plots for (T-bet, GATA3) axes, (T-bet, RORγT) axes and (GATA3, RORγT) axes. Here, we observed the hybrid ‘double positive’ states at a relatively higher frequency as compared to the toggle triad (**Fig 9D, S17, S18**). Hence, using the toggle triad model, we can predict the existence and provide a possible mechanistic explanation for the existence of stable hybrid helper T cell phenotypes, which has been shown experimentally at least for Th1/Th2 and Th1/Th17 cells.

## Discussion

Dissecting the dynamics of regulatory networks driving cellular differentiation and reprogramming is important to identify the trajectories that cells can take in the high-dimensional gene expression landscape as they commit to a cell-fate. Recent deluge in experimental technologies has enabled inferring these networks and identifying ‘master regulators’ of different cell-fates. Probing these networks from a dynamical systems perspective has helped characterize the ‘landscape’ of cell differentiation as proposed by Waddington over seven decades ago in which a cell – represented by a ball – rolls down into one or more possible branching valleys, each of which represents a stable cellular identity [29].

A frequent occurring network motif that has been identified in developmental decision-making and investigated from a dynamical systems perspective is a toggle switch – a double negative (hence, overall positive) feedback loop between two master regulators A and B. It has been shown to exhibit bistability with the two states being (high A, low B), and (low A, high B), representing a competition between A and B in enforcing the identity they drive and simultaneously repressing the one driven by their competitor [17]. Another well-studied motifs are negative feedback loops with two components (A activates B, B reduces the levels of A, such as p53-MDM2 [30]) or three components (A inhibits B, B inhibits C, and C inhibits A – a repressilator [20,31]) that can lead to sustained or damped oscillations. Often, such positive and negative feedback loops are intricately interlinked in natural biological networks to allow for a more diverse dynamic repertoire enabling bistability and/or oscillations [32–39]. Moreover, such feedback loops may be synchronized within a cell or across cellular populations to facilitate coordinated dynamics [40–42]. However, most theoretical attempts to investigate the coupled networks have been focused on bistable systems.

Here, we present a fundamentally different dynamical trait that can be achieved by three coupled toggle switches, or in other words, three mutually inhibitory ‘master regulators’ A, B and C forming a toggle triad. Our simulations show that the toggle triad network topology can enable tristability, with the three stable states being (high A, low B, low C), (low A, high B, low C), and (low A, low B, high C). Further, three more intermediate/hybrid states among these ‘single positive’ states can be enabled by a toggle triad, particularly when A, B and C can self-activate – (high A, high B, low C), (high A, low B, high C), (low A, high B, high C). In the Waddington landscape perspective, these three ‘double positive’ states can lie between two terminal ‘single positive’ states and can promote trans-differentiation among them (**Fig 10**). A previous modeling effort also suggested these possible six states for a toggle triad [16]; our approach through RACIPE enables two additional insights: a) a higher relative frequency of ‘single positive’ states as compared to the ‘double positive’ ones, and b) increase in ‘double positive’ states in presence of self-activation.

**Figure 10:**
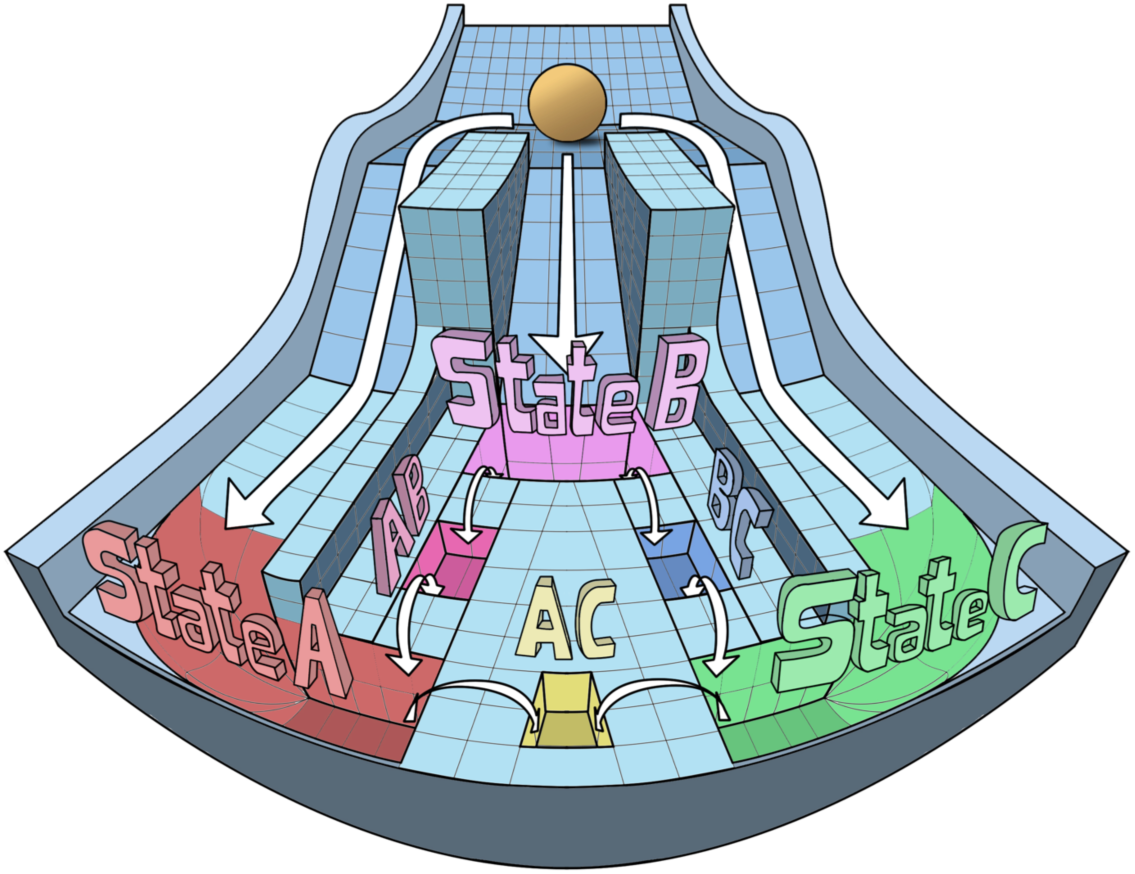
Waddington landscape for a toggle triad. Modified Waddington’s landscape to demonstrate the differentiation of three distinct ‘single positive’ states (states A, B and C), and three putative ‘double positive’ states (hybrid states A/B, A/C and C/B) from a common progenitor. These six states can be obtained from a toggle triad with/without self-activation.

We applied these results to reproduce the dynamics of naïve CD4^+^ helper T cells that differentiate to Th1, Th2, or Th17 [43]. Transcription factors T-bet, GATA3 and RORγT are considered to be the ‘master regulators’ of these cell fates respectively [28]. GATA3 and T-bet can self-activate directly or indirectly, and repress the activities or targets of one another [28,44], similar to a ‘toggle switch’ between two ‘master regulators’ seen in multiple scenarios [2]. In addition to the two mutually exclusive states (high GATA3, low T-bet and low GATA3, high T-bet in this case), self-activation can allow for the stable existence of a hybrid state (high GATA3, high T-bet in this case) [45–47]. Indeed, *in vitro* and *in vivo* experimental evidence has identified such ‘double positive’ individual cells stably co-expressing GATA3 and T-bet, referred to as a Th1/Th2 hybrid phenotype for the duration of weeks *in vitro* and months *in vivo* [46,48–51]. Intriguingly, hybrid Th1/Th2 cells can arise directly from the CD4+ naïve T cell precursors and/or reprogrammed from the Th2 cells [48,49], thus indicating phenotypic plasticity, a direct consequence of multistability in underlying biological networks [52]. Such plasticity can be explained by stochastic switching seen among the multiple states seen in a toggle triad with self-activation. Further, hybrid Th1/Th17 cells that stably persist *in vivo* have been experimentally identified; they express intermediate levels of both Th1 and Th17 signature transcription factors, and exhibit unique transcriptional and metabolic state as compared to Th1 and Th17 cells [53]. The stable *in vivo* existence of both hybrid Th1/Th2 and Th1/Th17 phenotypes suggests that their relationship may be expressed as ‘toggle switches’ with self-activation.

Recent single-cell analysis has strengthened the evidence for these hybrid phenotypes [54,55], revealing that the expression levels of various ‘master regulators’ in a population of cells is not limited to extreme high or low values [56], reminiscent of hybrid phenotypes seen in other biological scenarios [57,58]. Our analysis here is restricted to investigating steady-state (long-term) behaviors; however, approaches focusing on short-term dynamics of three-node and four-node networks have revealed important design principles related to noise attenuation [59–61]. The possible role of such functional traits in CD4+ T-cell decision-making needs to be investigated.

In addition to self-activation of T-bet and GATA3 discussed above, our model suggests self-activation of RORγT, which has been recently reported [27]. As an extension of this analysis, we propose a toggle triad involving Th1, Th2 and Th17 cells that reproduces all of the aforementioned predictions, and suggests that a hybrid Th2/Th17 cell could exist. While the stable existence of such cells and any phenotypic plasticity between Th2 and Th17 cells have not yet been reported, our model predicts their possible existence, especially when GATA3, RORγT and T-bet can self-activate themselves. Finally, the toggle triad model strengthens the hypothesis that the mixed cellular phenotypes are stable cellular identities with specific functional traits, and not just a transient co-expression of these lineage-determining transcription factors, as seen often in common progenitor cells [62].

Besides offering valuable insights into the dynamics of cellular decision-making, our results also pave the way towards designing tristable systems synthetically. Major efforts in synthetic biology have been, so far, targeted towards switches, cascades, pulse generators and oscillators [63–67]. The proposed network topology can be constructed to enable three distinct cell states, whereas including self-activation can facilitate the programming to achieve three hybrid cell states as well.

## Materials and Methods

### RACIPE (RAndom CIrcuit Perturbation) Analysis

#### Simulation

RACIPE is a computational tool that investigates the dynamics of a given network topology [22]. A topology file is given as the input to the program. It then simulates, for every parameter set, the network as a system of ordinary differential equations (ODEs) developed based on the input file. For every run of simulation, every kinetic parameter in the mathematical model is sampled from the defined biologically relevant range, thus giving multiple parameter sets. For each parameter set, the ODEs are solved for 100 initial conditions (default choice). The RACIPE simulation reports the steady state values for each component of the network for every parameter set in the solution file. For all our analyses, we have used the default ranges for sampling the parameters, sampled 10000 parameter sets and 1000 initial conditions for every parameter set. Depending on the number of steady states the system converges to for a given parameter set, the system is classified as monostable, bistable, tristable all the way to decastable. Even if one initial condition converges to a different steady state, RACIPE considers the system to be multistable for that parameter set.

A generic differential equation for component A affected by component B is denoted by RACIPE as:

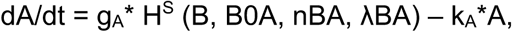

where g_A_ and k_A_ are corresponding production and degradation rates, H^S^ (B, B0A, nBA, λBA) denotes a shifted Hill function defined as:

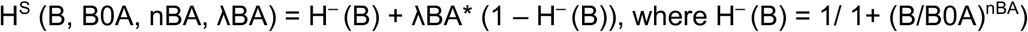

#### Normalization of Steady states

The steady state solution provided by RACIPE simulation are in log2 scale. We normalized the obtained steady states in the solution files two-fold. To account for extremes in sampling of the production and degradation rate parameters, we performed g/k normalization. We divided every steady state value (E_i_) in the solution file by the ratio of the production and degradation rate of the respective component (g_i_/k_i_) of the network of the corresponding parameter set. Following that, we performed z-score normalization. We calculated the mean (E_in_) and standard deviation (σ_in_) for every component ‘i’ over all parameter sets after the g/k normalization. The final transformation formula for every steady state is as follows:

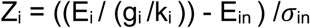

where Z_i_ is the final normalized expression. We found the distributions of every component to be largely bimodal in nature (Fig S2) with the center of the two modes to be around 0. Thus, we chose to define the states “high” and “low” as greater than and smaller than 0 respectively.

#### Clustering and Replicates

For the hierarchical clustering shown in heatmaps (Fig 3, 5) and other supplementary figures (Fig S4-S9), the clustergram function in MATLAB was used. For coloring of the scatter plots in Fig 6, k-means clustering (k = 6) was used to identify the clusters. Since k-means clustering can provide variant results for every run of the function, we confirmed the clusters by running the clustering function for the same data thrice, the latter two replicates (Fig S17, S18) for the networks of Toggle Triad (TT) and Toggle triad with 3 self-activations (TT+3SA) respectively. For every network shown in main text and SI, RACIPE simulations were run thrice to obtain three independent replicates. Further analyses was then performed on these replicates, with the data presented as mean +/- standard deviation (shown by error bars).

### Boolean Analysis

For Boolean analysis as well for TT and C2 circuits, a topological file is given as the input. The file determines the nodes and edges of the network. The edges are of two types, activatory and inhibitory. The analyses carried out were by two methods, synchronous and asynchronous update of the nodes. The constraint of equal weightage to inhibitory and activating links was used [23,68]. The updating of the nodes follows a simple majority rule. The node is updated to 1 if the sum of activations to the node is higher than inhibitions and updated to 0 for the opposite case. The steady state is said to be reached if there is no change in the updates for a few time-steps. We have run the simulations for 10000 initial conditions randomly sampled over all possible states of the network.

### Bifurcation Analysis

Bifurcation diagrams were plotted using the continuation software PyDSTool [69].

### sRACIPE simulations

We performed sRACIPE simulations on either the TT or TT+3SA to generate a set of random parameter sets and simulated the system with a fixed amount of noise in one of the parameters. We used the webserver facility of Gene Circuit Explorer (GeneEx) to simulate stochastic dynamics of gene regulatory circuits - https://shinyapps.jax.org/5c965c4b284ca029b4aa98483f3da3c5/

## Acknowledgements

This work was supported by Ramanujan Fellowship (SB/S2/RJN-049/2018) awarded to MKJ by Science and Engineering Research Board, Department of Science and Technology, Government of India.

## Author contributions

MKJ designed and supervised research; ASD, SS and SH conducted research; ASD, SS, SJ, MKJ analyzed data; all authors participated in writing and revision of the manuscript.

## References

1. Guantes R, Poyatos JF. 2008 Multistable decision switches for flexible control of epigenetic differentiation. PLoS Comput. Biol. 4, e1000235. (doi: 10.1371/journal.pcbi.1000235)

2. Zhou JX, Huang S. 2011 Understanding gene circuits at cell-fate branch points for rational cell reprogramming. Trends Genet. 27, 55–62. (doi: 10.1016/j.tig.2010.11.002)

3. Qian Y, McBride C, Del Vecchio D. 2018 Programming Cells to Work for Us. Annu. Rev. Control. Robot. Auton. Syst. 1, 411–40. (doi: 10.1146/annurev-control-060117-105052)

4. Chang HH, Oh PY, Ingber DE, Huang S. 2006 Multistable and multistep dynamics in neutrophil differentiation. BMC Cell Biol. 7, 11. (doi: 10.1186/1471-2121-7-11)

5. Novick A, Weiner M. 1957 Enzyme induction as an all-or-none phenomenon. Proc. Natl. Acad. Sci. U. S. A. 43, 553–66. (doi: 10.1073/pnas.43.7.553)

6. Ferrell JE. 2012 Bistability, Bifurcations, and Waddington’s Epigenetic Landscape. Curr. Biol. 22, R458–R466. (doi: 10.1016/J.CUB.2012.03.045)

7. Gardner TS, Cantor CR, Collins JJ. 2000 Construction of a genetic toggle switch in Escherichia coli. Nature 403, 339–342. (doi: 10.1038/35002131)

8. Veening J, Smits WK, Kuipers OP. 2008 Bistability, Epigenetics, and Bet-Hedging in Bacteria. Annu. Rev. Microbiol. 62, 193–212. (doi: 10.1146/annurev.micro.62.081307.163002)

9. Ozbudak EM, Thattai M, Lim HN, Shraiman BI, van Oudenaarden A. 2004 Multistability in the lactose utilization network of Escherichia coli. Nature 427, 737–40. (doi: 10.1038/nature02298)

10. Celià-Terrassa T et al. 2018 Hysteresis control of epithelial-mesenchymal transition dynamics conveys a distinct program with enhanced metastatic ability. Nat. Commun. 9, 5005. (doi: 10.1038/s41467-018-07538-7)

11. Jaruszewicz J, Lipniacki T. 2013 Toggle switch: noise determines the winning gene. Phys. Biol. 10, 035007. (doi: 10.1088/1478-3975/10/3/035007)

12. Ma R, Wang J, Hou Z, Liu H. 2012 Small-number effects: A third stable state in a genetic bistable toggle switch. Phys. Rev. Lett. 109, 248107. (doi: 10.1103/PhysRevLett.109.248107)

13. Tian T, Burrage K. 2006 Stochastic models for regulatory networks of the genetic toggle switch. Proc. Natl. Acad. Sci. U. S. A. 103, 8372–8377. (doi: 10.1073/pnas.0507818103)

14. Lipshtat A, Loinger A, Balaban NQ, Biham O. 2006 Genetic toggle switch without cooperative binding. Phys. Rev. Lett. 96, 188101. (doi: 10.1103/PhysRevLett.96.188101)

15. Jia D, Jolly MK, Harrison W, Boareto M, Ben-Jacob E, Levine H. 2017 Operating principles of tristable circuits regulating cellular differentiation. Phys. Biol. 14, 035007.

16. Leon M, Woods ML, Fedorec AJH, Barnes CP. 2016 A computational method for the investigation of multistable systems and its application to genetic switches. BMC Syst. Biol. 10, 130.

17. Shea JJO, Paul WE, Cells CDT. 2012 Mechanisms underlying linear commitment and plasticity of helper CD4+ T cells. Science (80-.). 1098, 1098–1103. (doi: 10.1126/science.1178334)

18. Fang D, Zhu J. 2017 Dynamic balance between master transcription factors determines the fates and functions of CD4 T cell and innate lymphoid cell subsets. J. Exp. Med. 214, 1861–1876. (doi: 10.1084/jem.20170494)

19. Ling G, Guan Z-H, He D-X, Liao R-Q, Zhang X-H. 2014 Stability and bifurcation analysis of new coupled repressilators in genetic regulatory networks with delays. Neural Netw. 60, 222–231. (doi: 10.1016/j.neunet.2014.08.012)

20. Elowitz MB, Leibier S. 2000 A synthetic oscillatory network of transcriptional regulators. Nature 400, 335–338. (doi: 10.1038/35002125)

21. Müller S, Hofbauer J, Endler L, Flamm C, Widder S, Schuster P. 2006 A generalized model of the repressilator. J. Math. Biol. 53, 905–937. (doi: 10.1007/s00285-006-0035-9)

22. Huang B, Lu M, Jia D, Ben-Jacob E, Levine H, Onuchic JN. 2017 Interrogating the topological robustness of gene regulatory circuits by randomization. PLoS Comput. Biol. 13, e1005456. (doi: 10.137/journal.pcbi.1005456)

23. Font-Clos F, Zapperi S, La Porta CAM. 2018 Topography of epithelial–mesenchymal plasticity. Proc. Natl. Acad. Sci. 115, 5902–5907. (doi: 10.1073/pnas.1722609115)

24. Thomas P, Popovic N, Grima R. 2014 Phenotypic switching in gene regulatory networks. Proc Natl Acad Sci U S A 111, 6994–6999. (doi: 10.1073/pnas.1400049111)

25. Kohar V, Lu M. 2018 Role of noise and parametric variation in the dynamics of gene regulatory circuits. npj Syst. Biol. Appl. 4, 40. (doi: 10.1038/s41540-018-0076-x)

26. Evans CM, Jenner RG. 2013 Transcription factor interplay in t helper cell differentiation. Brief. Funct. Genomics 12, 499–511. (doi: 10.1093/bfgp/elt025)

27. Xiao S et al. 2014 Small-molecule RORγt antagonists inhibit T helper 17 cell transcriptional network by divergent mechanisms. Immunity 40, 477–489. (doi: 10.1016/j.immuni.2014.04.004)

28. Murphy KM, Stockinger B. 2010 Effector T cell plasticity: Flexibility in the face of changing circumstances. Nat. Immunol. 11, 674–680. (doi: 10.1038/ni.1899)

29. Furusawa C, Kaneko K. 2012 A dynamical-systems view of stem cell biology. Science (80-.). 338, 215–217. (doi: 10.1126/science.1224311)

30. Bar-Or RL, Maya R, Segel LA, Alon U, Levine AJ, Oren M. 2000 Generation of oscillations by the p53-Mdm2 feedback loop: A theoretical and experimental study. Proc. Natl. Acad. Sci. U. S. A. 97, 11250–11255. (doi: 10.1073/pnas.210171597)

31. Buse O, Pérez R, Kuznetsov A. 2010 Dynamical properties of the repressilator model. Phys. Rev. E - Stat. Nonlinear, Soft Matter Phys. 81, 066206. (doi: 10.1103/PhysRevE.81.066206)

32. Tian XJ, Zhang XP, Liu F, Wang W. 2009 Interlinking positive and negative feedback loops creates a tunable motif in gene regulatory networks. Phys. Rev. E - Stat. Nonlinear, Soft Matter Phys. 80, 1–8. (doi: 10.1103/PhysRevE.80.011926)

33. Tiwari A, Igoshin OA. 2012 Coupling between feedback loops in autoregulatory networks affects bistability range, open-loop gain and switching times. Phys. Biol. 9, 055003. (doi: 10.1088/1478-3975/9/5/055003)

34. Pfeuty B, Kaneko K. 2009 The combination of positive and negative feedback loops confers exquisite flexibility to biochemical switches. Phys. Biol. 6, 046013. (doi: 10.1088/1478-3975/6/4/046013)

35. Avendaño MS, Leidy C, Pedraza JM. 2013 Tuning the range and stability of multiple phenotypic states with coupled positive–negative feedback loops. Nat. Commun. 4, 2605. (doi: 10.1038/ncomms3605)

36. Krishna S, Semsey S, Jensen MH. 2009 Frustrated bistability as a means to engineer oscillations in biological systems. Phys. Biol. 6, 036009. (doi: 10.1088/1478-3975/6/3/036009)

37. Wang LS, Li NX, Chen JJ, Zhang XP, Liu F, Wang W. 2018 Modulation of dynamic modes by interplay between positive and negative feedback loops in gene regulatory networks. Phys. Rev. E 97, 042412. (doi: 10.1103/PhysRevE.97.042412)

38. Page KM. 2019 Oscillations in well-mixed, deterministic feedback systems: Beyond ring oscillators. J. Theor. Biol. 481, 44–53. (doi: 10.1016/j.jtbi.2019.05.004)

39. Perez-Carrasco R, Barnes CP, Schaerli Y, Isalan M, Briscoe J, Page KM. 2018 Combining a Toggle Switch and a Repressilator within the AC-DC Circuit Generates Distinct Dynamical Behaviors. Cell Syst. 6, 521–530. (doi: 10.1016/j.cels.2018.02.008)

40. Nguyen C, Han SK. 2010 Synchronization of toggle switches coupled through a common inhibitor. Europhys. Lett. 90, 10010. (doi: 10.1209/0295-5075/90/10010)

41. Fernandez-Niño M, Giraldo D, Gomez-Porras JL, Dreyer I, Barrios AFG, Arevalo-Ferro C. 2017 A synthetic multi-cellular network of coupled self-sustained oscillators. PLoS One 12, e0180155. (doi: 10.1371/journal.pone.0180155)

42. Hellen EH, Volkov E. 2017 Flexible dynamics of two quorum-sensing coupled repressilators. Phys. Rev. E 95, 022408. (doi: 10.1103/PhysRevE.95.022408)

43. Zhu J, Yamane H, Paul WE. 2010 Differentiation of Effector CD4 T Cell Populations. Annu. Rev. Immunol. 28, 445–89. (doi: 10.1146/annurev-immunol-030409-101212)

44. Geginat J, Paroni M, Maglie S, Alfen JS, Kastirr I, Gruarin P, de Simone M, Pagani M, Abrignani S. 2014 Plasticity of human CD4 T cell subsets. Front. Immunol. 5, 630. (doi: 10.3389/fimmu.2014.00630)

45. Jolly MK, Boareto M, Lu M, Onuchic JN, Clementi C, Ben-Jacob E. 2015 Operating principles of Notch-Delta-Jagged module of cell-cell communication. New J. Phys. 17, 55021.

46. Antebi YE, Reich-Zeliger S, Hart Y, Mayo A, Eizenberg I, Rimer J, Putheti P, Pe’er D, Friedman N. 2013 Mapping differentiation under mixed culture conditions reveals a tunable continuum of T cell fates. PLoS Biol. 11, e1001616. (doi: 10.1371/journal.pbio.1001616)

47. Yates A, Callard R, Stark J. 2004 Combining cytokine signalling with T-bet and GATA-3 regulation in Th1 and Th2 differentiation: A model for cellular decision-making. J. Theor. Biol. 231, 181–96. (doi: 10.1016/j.jtbi.2004.06.013)

48. Peine M et al. 2013 Stable T-bet+GATA-3+ Th1/Th2 Hybrid Cells Arise In Vivo, Can Develop Directly from Naive Precursors, and Limit Immunopathologic Inflammation. PLoS Biol. 11, e1001633. (doi: 10.1371/journal.pbio.1001633)

49. Hegazy AN et al. 2010 Interferons Direct Th2 Cell Reprogramming to Generate a Stable GATA-3+T-bet+ Cell Subset with Combined Th2 and Th1 Cell Functions. Immunity 32, 116–128. (doi: 10.1016/j.immuni.2009.12.004)

50. Huang S. 2013 Hybrid T-Helper Cells: Stabilizing the Moderate Center in a Polarized System. PLoS Biol. 11, e1001632. (doi: 10.1371/journal.pbio.1001632)

51. Fang M, Xie H, Dougan SK, Ploegh H. 2013 Stochastic Cytokine Expression Induces Mixed T Helper Cell States. PLoS Biol. 11, e1001618. (doi: 10.1371/journal.pbio.1001618)

52. Jia D, Jolly MK, Kulkarni P, Levine H. 2017 Phenotypic Plasticity and Cell Fate Decisions in Cancer : Insights from Dynamical Systems Theory. Cancers (Basel). 9, E70. (doi: 10.3390/cancers9070070)

53. Chatterjee S et al. 2018 CD38-NAD + Axis Regulates Immunotherapeutic Anti-Tumor T Cell Response. Cell Metab. 27, 85-100.e8. (doi: 10.1016/j.cmet.2017.10.006)

54. Xhangholi I, Dura B, Lee G, Kim D, Xiao Y, Fan R. 2019 Single-cell Analysis of CAR-T Cell Activation Reveals A Mixed TH1/TH2 Response Independent of Differentiation. Genomics. Proteomics Bioinformatics 17, 129–139.

55. MacLean AL, Hong T, Nie Q. 2018 Exploring intermediate cell states through the lens of single cells. Curr. Opin. Syst. Biol. 9, 32–41. (doi: 10.1016/j.coisb.2018.02.009)

56. Eizenberg-Magar I, Rimer J, Zaretsky I, Lara-Astiaso D, Reich-Zeliger S, Friedman N. 2017 Diverse continuum of CD4+ T-cell states is determined by hierarchical additive integration of cytokine signals. Proc. Natl. Acad. Sci. U. S. A. 114, E6647–E6456. (doi: 10.1073/pnas.1615590114)

57. Jolly MK, Mani SA, Levine H. 2018 Hybrid epithelial/mesenchymal phenotype(s): The ‘fittest’ for metastasis? Biochim. Biophys. Acta - Rev. Cancer 1870, 151–157. (doi: 10.1016/j.bbcan.2018.07.001)

58. Huang B, Lu M, Jolly MK, Tsarfaty I, Onuchic J, Ben-Jacob E. 2014 The three-way switch operation of Rac1/RhoA GTPase-based circuit controlling amoeboid-hybrid-mesenchymal transition. Sci. Rep. 4, 6449. (doi: 10.1038/srep06449)

59. Ma W, Trusina A, El-Samad H, Lim WA, Tang C. 2009 Defining Network Topologies that Can Achieve Biochemical Adaptation. Cell 138, 76–73. (doi: 10.1016/j.cell.2009.06.013)

60. Chen M, Wang L, Liu CC, Nie Q. 2013 Noise attenuation in the on and off states of biological switches. ACS Synth. Biol. 2, 587–593. (doi: 10.1021/sb400044g)

61. Qiao L, Zhao W, Tang C, Nie Q, Zhang L. 2019 Network Topologies That Can Achieve Dual Function of Adaptation and Noise Attenuation. Cell Syst. 9, 271–285. (doi: 10.1016/j.cels.2019.08.006)

62. Oestreich KJ, Weinmann AS. 2012 Master regulators or lineage-specifying? Changing views on CD4+ T cell transcription factors. Nat. Rev. Immunol. 12, 799–804. (doi: 10.1038/nri3321)

63. Tigges M, Marquez-Lago TT, Stelling J, Fussenegger M. 2009 A tunable synthetic mammalian oscillator. Nature 457, 309–312. (doi: 10.1038/nature07616)

64. Atkinson MR, Savageau MA, Myers JT, Ninfa AJ. 2003 Development of genetic circuitry exhibiting toggle switch or oscillatory behavior in Escherichia coli. Cell 113, 597–607. (doi: 10.1016/S0092-8674(03)00346-5)

65. Basu S, Mehreja R, Thiberge S, Chen MT, Weiss R. 2004 Spatiotemporal control of gene expression with pulse-generating networks. Proc. Natl. Acad. Sci. U. S. A. 101, 6355–6360. (doi: 10.1073/pnas.0307571101)

66. Kramer BP, Viretta AU, Baba MD El, Aubel D, Weber W, Fussenegger M. 2004 An engineered epigenetic transgene switch in mammalian cells. Nat. Biotechnol. 22, 867–870. (doi: 10.1038/nbt980)

67. Stricker J, Cookson S, Bennett MR, Mather WH, Tsimring LS, Hasty J. 2008 A fast, robust and tunable synthetic gene oscillator. Nature 456, 516–519. (doi: 10.1038/nature07389)

68. Hari K, Sabuwala B, Subramani BV, Porta C La, Zapperi S, Font-Clos F, Jolly MK. 2020 Identifying inhibitors of epithelial-mesenchymal plasticity using a network topology based approach. npj Syst. Biol. Appl. 6, 15. (doi: 10.1038/s41540-020-0132-1)

69. Clewley R. 2012 Hybrid models and biological model reduction with PyDSTool. PLoS Comput. Biol. 8, e1002628. (doi: 10.1371/journal.pcbi.1002628)

