## Supplementary Figures for "Multistability in cellular differentiation enabled by a network of three mutually repressing master regulators"

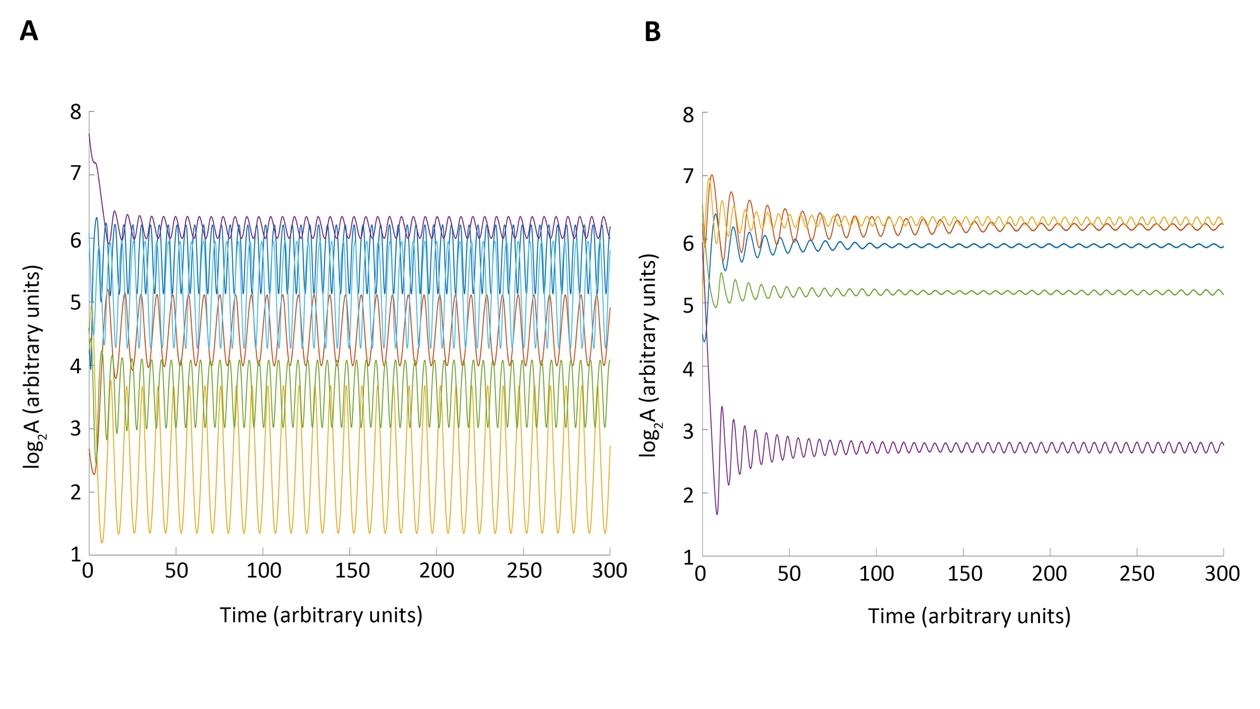

**Figure S1**: Oscillatory behavior of Repressilator. A, B) Representative cases for different parameter sets showing sustained or damped oscillations. Such parameter sets usually get erroneously binned into ‘decastable’ scenarios in RACIPE due to the software being unable to find a stable steady state for stipulated period of time.

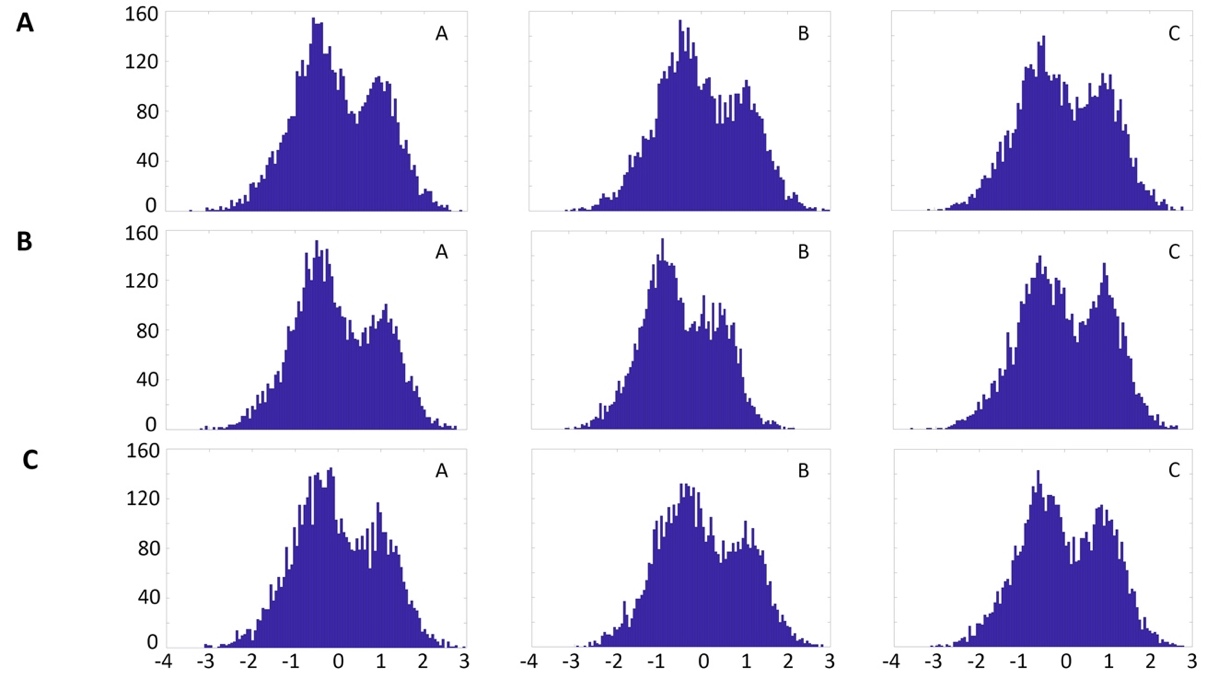

**Figure S2**: Bimodal distributions of the three components of toggle triad network. A) Distributions of normalized levels of A, B and C for all monostable solutions seen for the first RACIPE replicate. B, C) Same as A but for second and third RACIRE replicate. X-axis of the histograms represent the z-scores of the respective component of the corresponding replicate. Y-axis of the histograms represent the frequency.

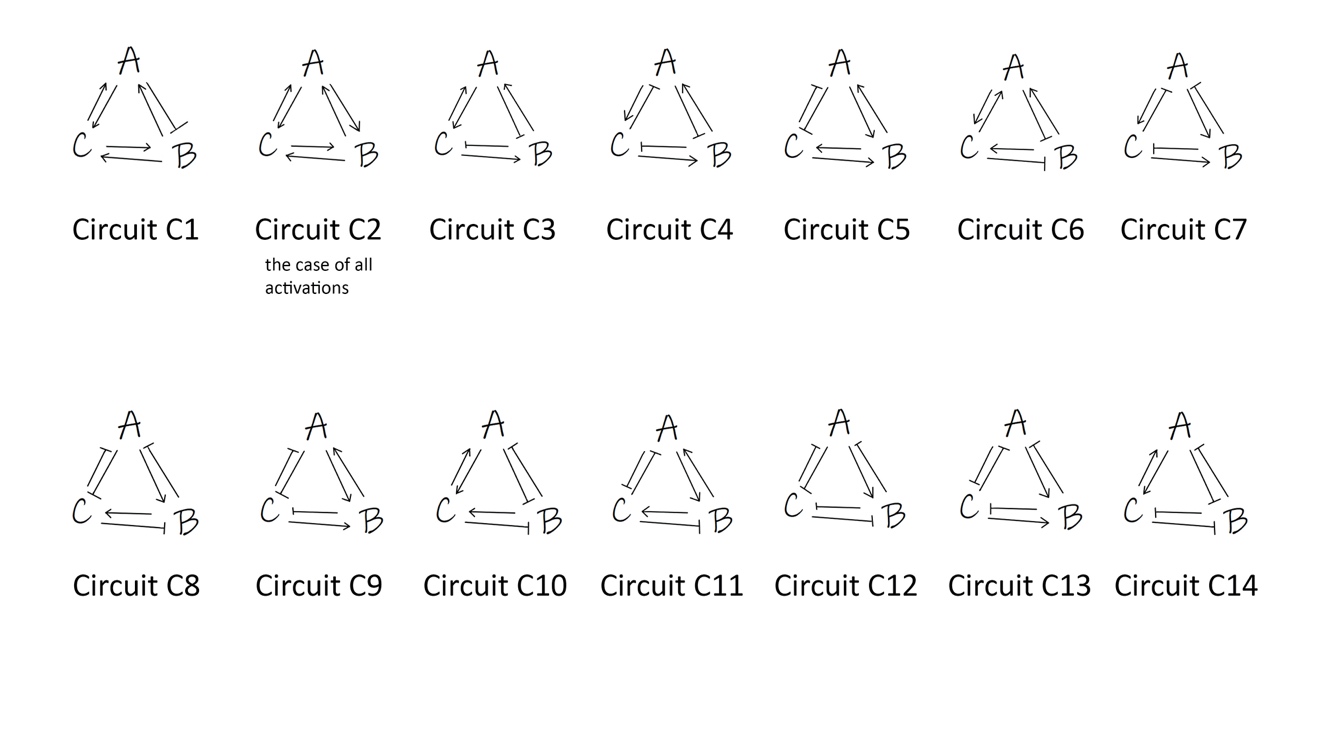

**Figure S3**: Three component network circuits with combinations of inhibitory links in the toggle triad replaced by activating links.

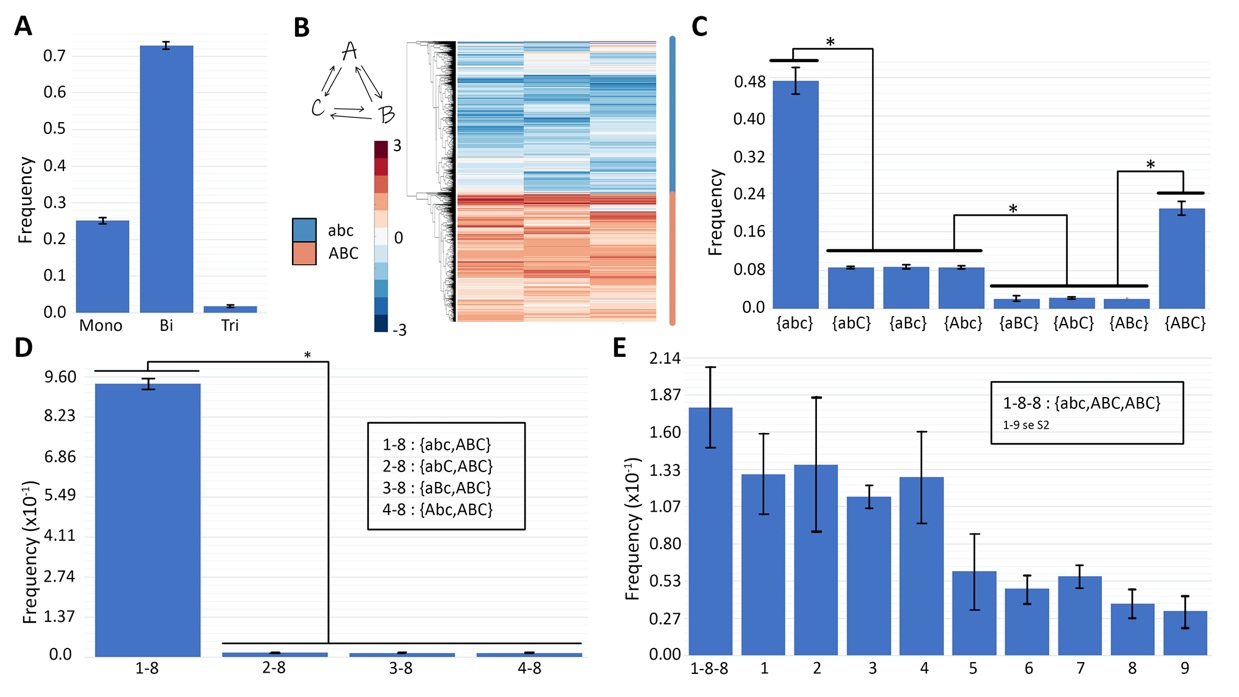

**Figure S4**: Characterization of circuit C2. A) Frequency of monostable, bistable and tristable solutions. B) Heatmap showing all solutions for circuit C2; the nomenclature shown capitalizes the node whose levels are relatively high – {abc} = (low A, low B, low C), {ABC} = (high A, high B, high C). C) Frequency of 8 monostable solutions. D, E) Frequencies for different bistable and tristable phase; most frequent bistable and tristable phases are combinations of abc and ABC.

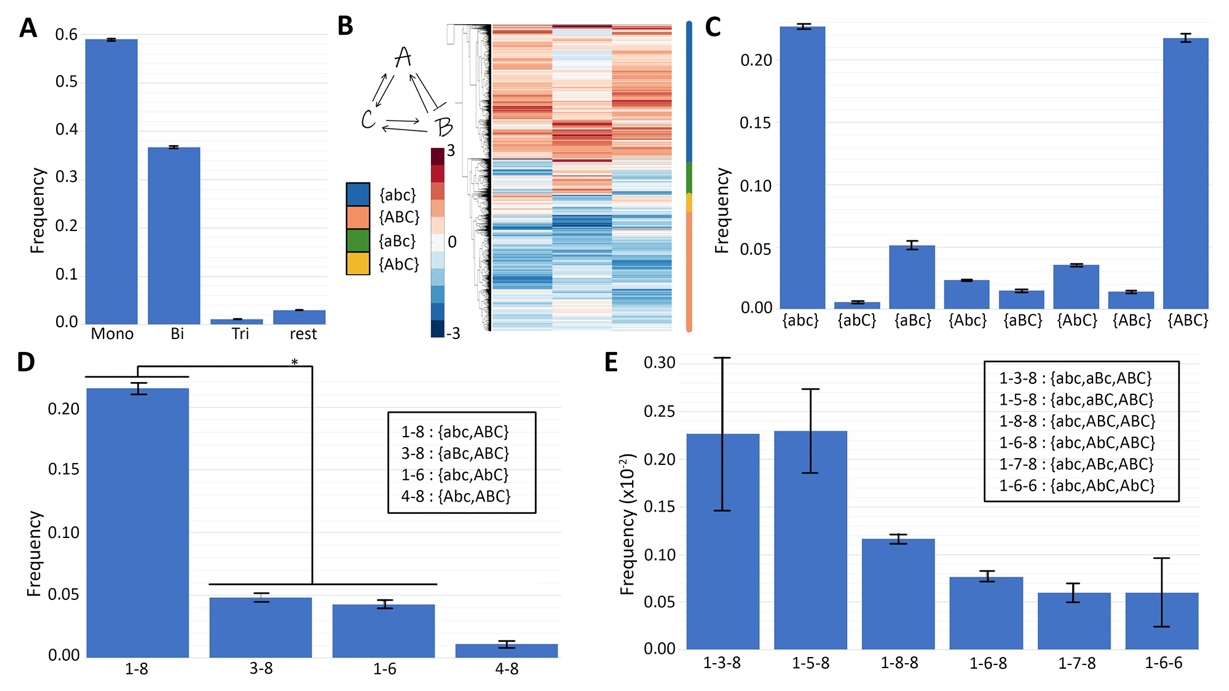

**Figure S5**: Characterization of circuit C1. A) Frequency of monostable, bistable, tristable solutions. B) Heatmap showing all solutions for circuit C1; the nomenclature shown capitalizes the node whose levels are relatively high – {abc}= (low A, low B, low C), {ABC} = (high A, high B, high C). C) Frequency of the 8 = (2^3) monostable solutions. D, E) Most frequent bistable and tristable phases are combinations of abc and ABC.

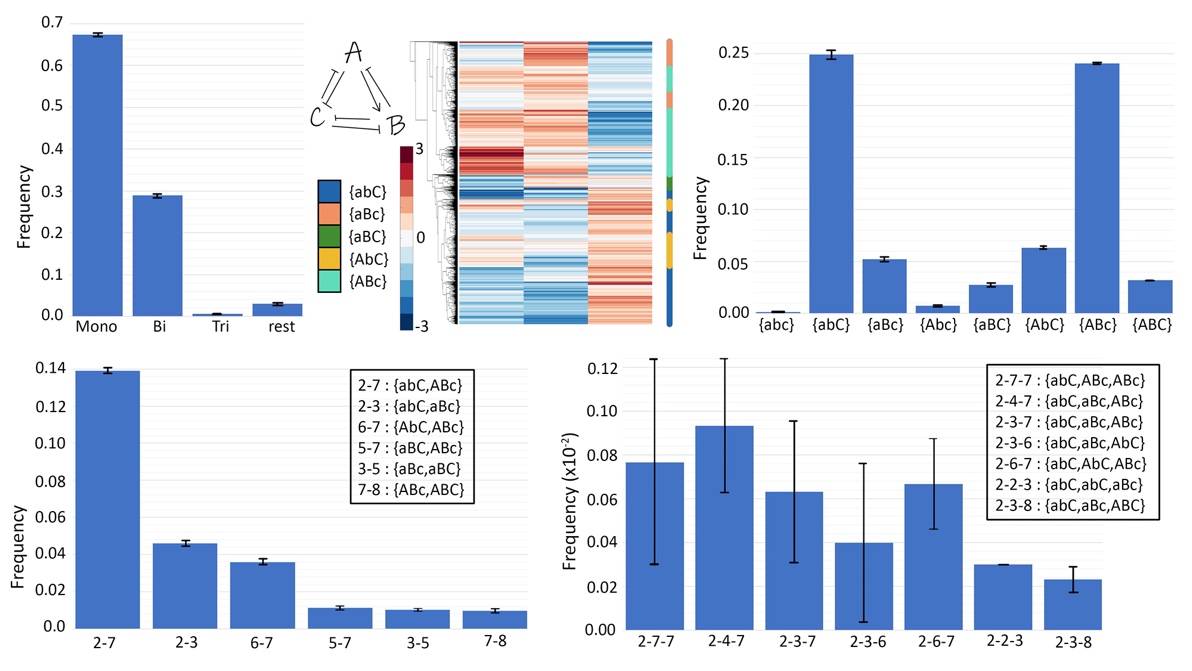

**Figure S6**: Characterization of circuit C12. A) Frequency of monostable, bistable, tristable solutions. B) Heatmap showing all solutions for circuit C12; the nomenclature shown capitalizes the node whose levels are relatively high – {Abc}= (A-high, Blow, C-low}, {aBc} = (A-low, B-high, C-low), {abC} = (A-low, B-low, C-high. C) Frequency of the 8 = (2^3) monostable solutions. D, E) Most frequent bistable and tristable phases are combinations of abC and ABc.

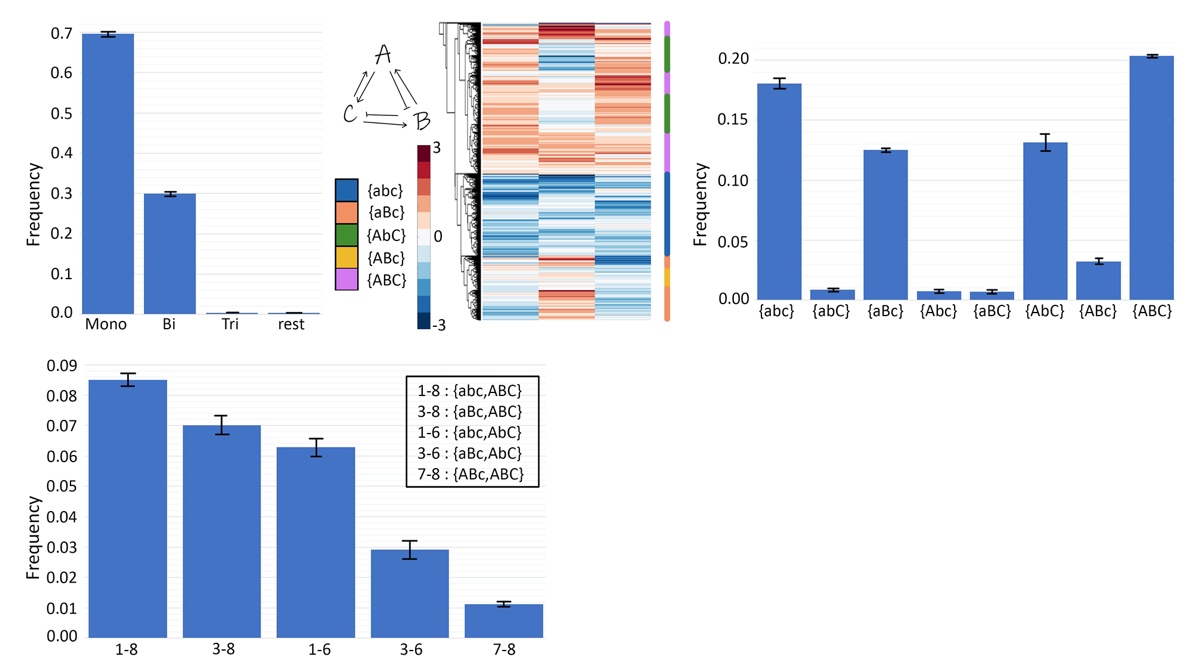

**Figure S7**: Characterization of circuit C3. A) Frequency of monostable, bistable, tristable solutions. B) Heatmap showing all solutions for circuit C3; the nomenclature shown capitalizes the node whose levels are relatively high – {abc}= (low A, low B, low C), {ABC} = (high A, high B, high C). C) Frequency of the 8 = (2^3) monostable solutions. D) Most frequent bistable phases are combinations of abc, ABC, aBc and AbC.

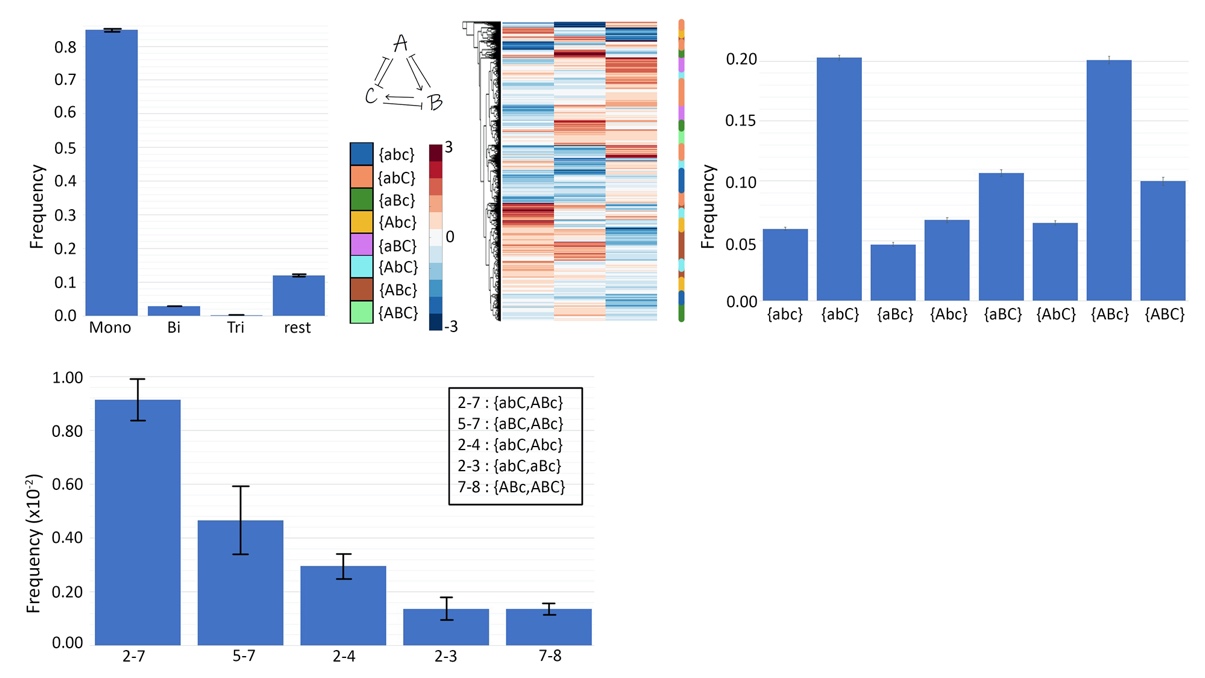

**Figure S8**: Characterization of circuit C8. A) Frequency of monostable, bistable, tristable solutions. B) Heatmap showing all solutions for circuit C8; the nomenclature shown capitalizes the node whose levels are relatively high – {Abc}= (high A, B low, C low}, {aBc} = (low A, high B, low C), {abC} = (low A, low B, high C). C) Frequency of the 8 = (2^3) monostable solutions. D) Most frequent bistable phases are combinations of abC, ABc and aBc.

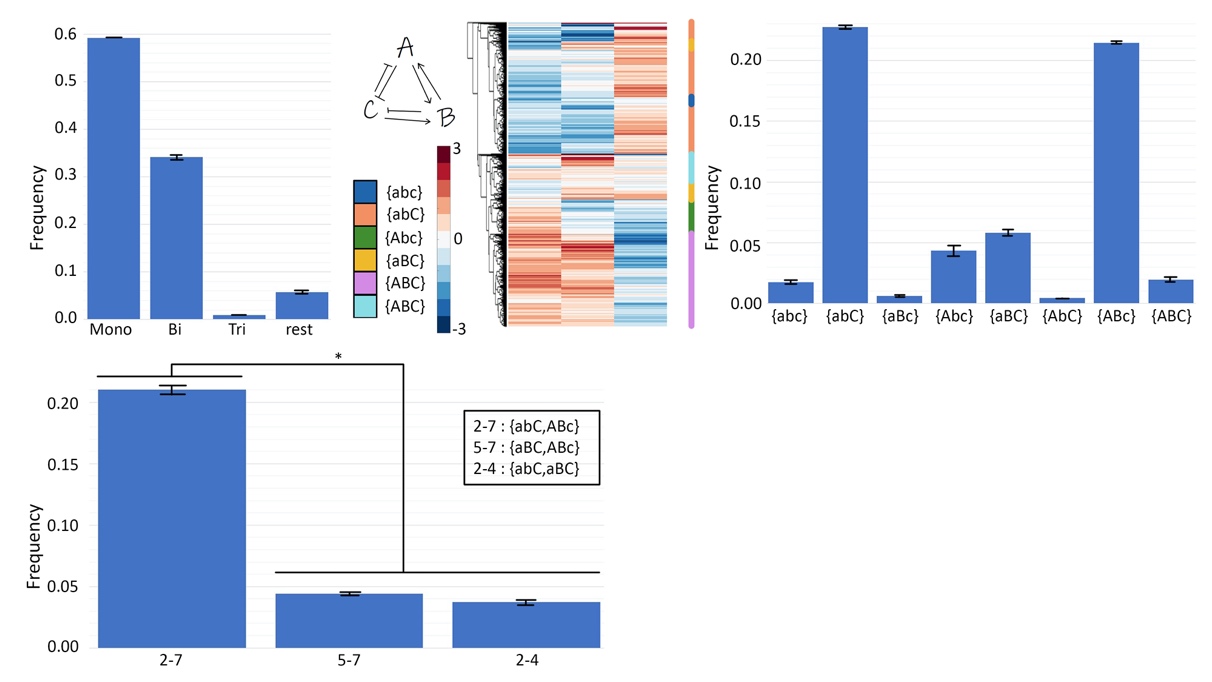

**Figure S9**: Characterization of circuit C9. A) Frequency of monostable, bistable, tristable solutions. B) Heatmap showing all solutions for circuit C9; the nomenclature shown capitalizes the node whose levels are relatively high – {ABc}= (high A, high B, low C}, {abC} = (low A, low B, high C). C) Frequency of the 8 = (2^3) monostable solutions. D, E) Most frequent bistable and tristable phases are combinations of abC and ABc.

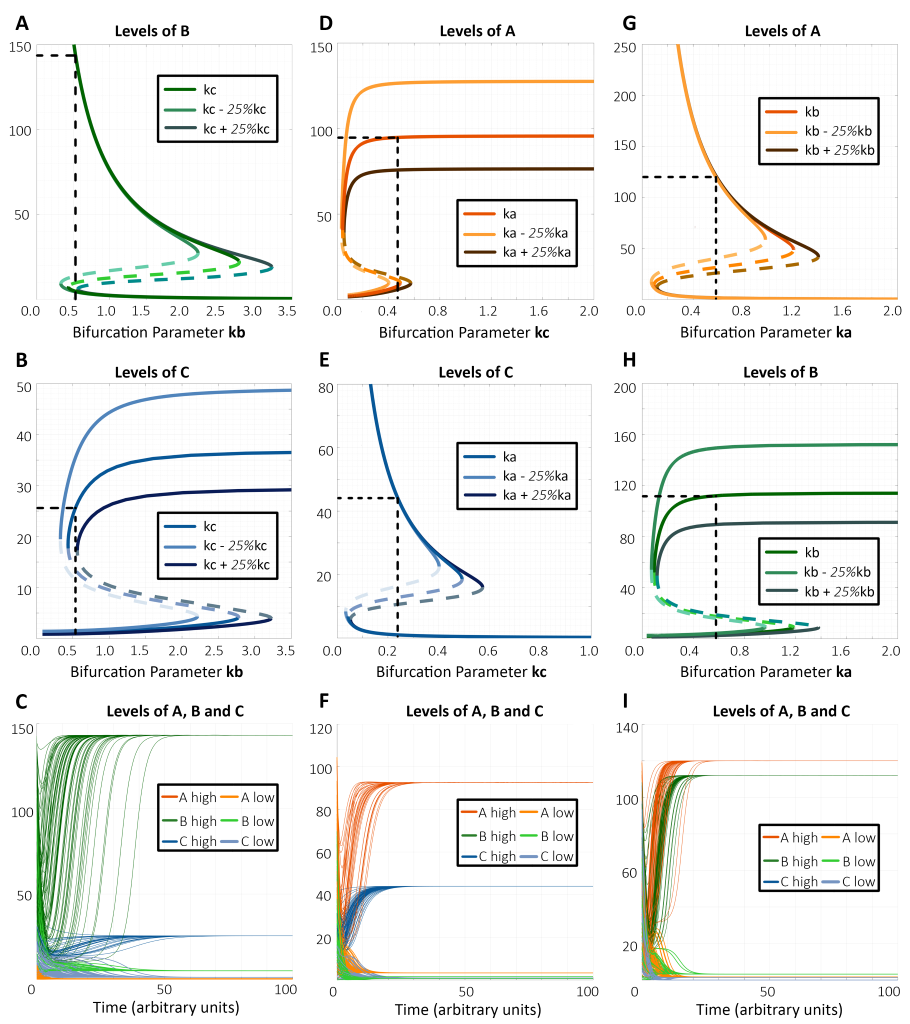

**Figure SI0**: Bifurcation diagrams and dynamics plots for representative cases of bistable phases. A) Bifurcation diagram of expression level of component B with kb as bifurcation parameter for the bistable phase {aBc, abC}. B) Same as B) but for component C. C) Dynamics plots of expression levels of components A, B and C for the bistable phase {aBc, abC}, showing convergence to two different states with varied levels of B and C (levels of A are low in both cases). D) Bifurcation diagram of expression level of component A with kc as the bifurcation parameter for the bistable phase {Abc, abC}. E) Same as D) but for component C. F) Dynamics plots of expression levels of components A, B and C for the bistable phase {Abc, abC}, showing convergence to two different levels of A and C (levels of B are low in both cases) G) Bifurcation diagram of expression level of component A with ka as bifurcation parameter for bistable phase {aBc, Abc}. H) Same as G) but for component B. I) Dynamics plots of expression levels of components A, B and C for bistable phase {aBc, Abc}, showing convergence to two different values of A and B (levels of C are low for both cases). The parameter values are given correspondingly in Table S13.

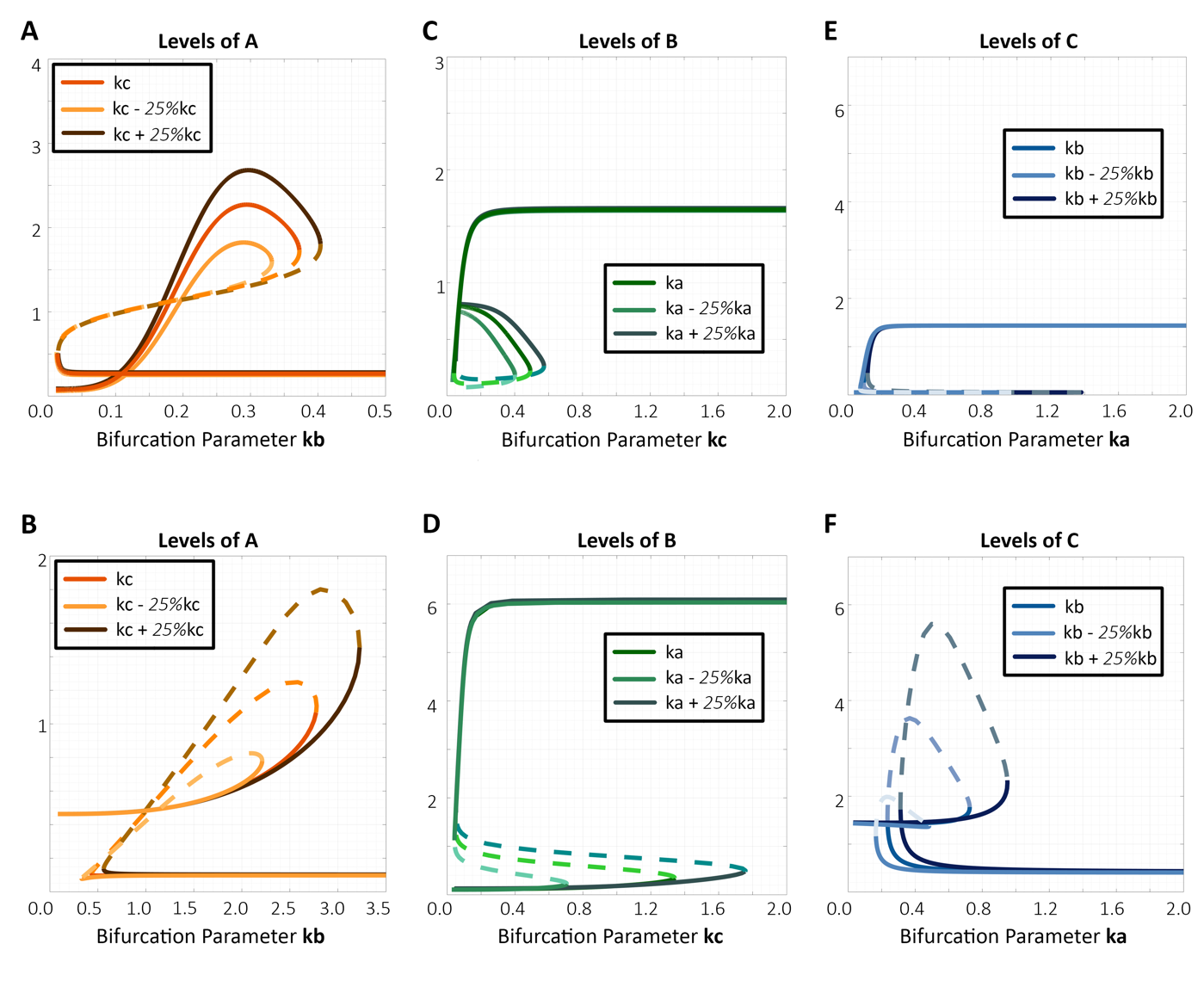

**Figure S11**: Bifurcation diagrams for the component expressing low levels in all the respective bistable phases’ representative cases shown in Fig 4 and Fig S10. A) Bifurcation diagram of expression level of component A with kb as bifurcation parameter for the bistable phase {aBc, abC} as in Fig 4. B) Bifurcation diagram of expression level of component A with kb as bifurcation parameter for the bistable phase {aBc, abC} as in Fig S10. C) Bifurcation diagram of expression level of component B with kc as bifurcation parameter for the bistable phase {Abc, abC} as in Fig 4. D) Bifurcation diagram of expression level of component B with kc as bifurcation parameter for the bistable phase {Abc, abC} as in Fig S10. E) Bifurcation diagram of expression level of component C with ka as bifurcation parameter for the bistable phase {aBc, Abc} as in Fig 4. F) Bifurcation diagram of expression level of component C with ka as bifurcation parameter for the bistable phase {aBc, Abc} as in Fig S10.

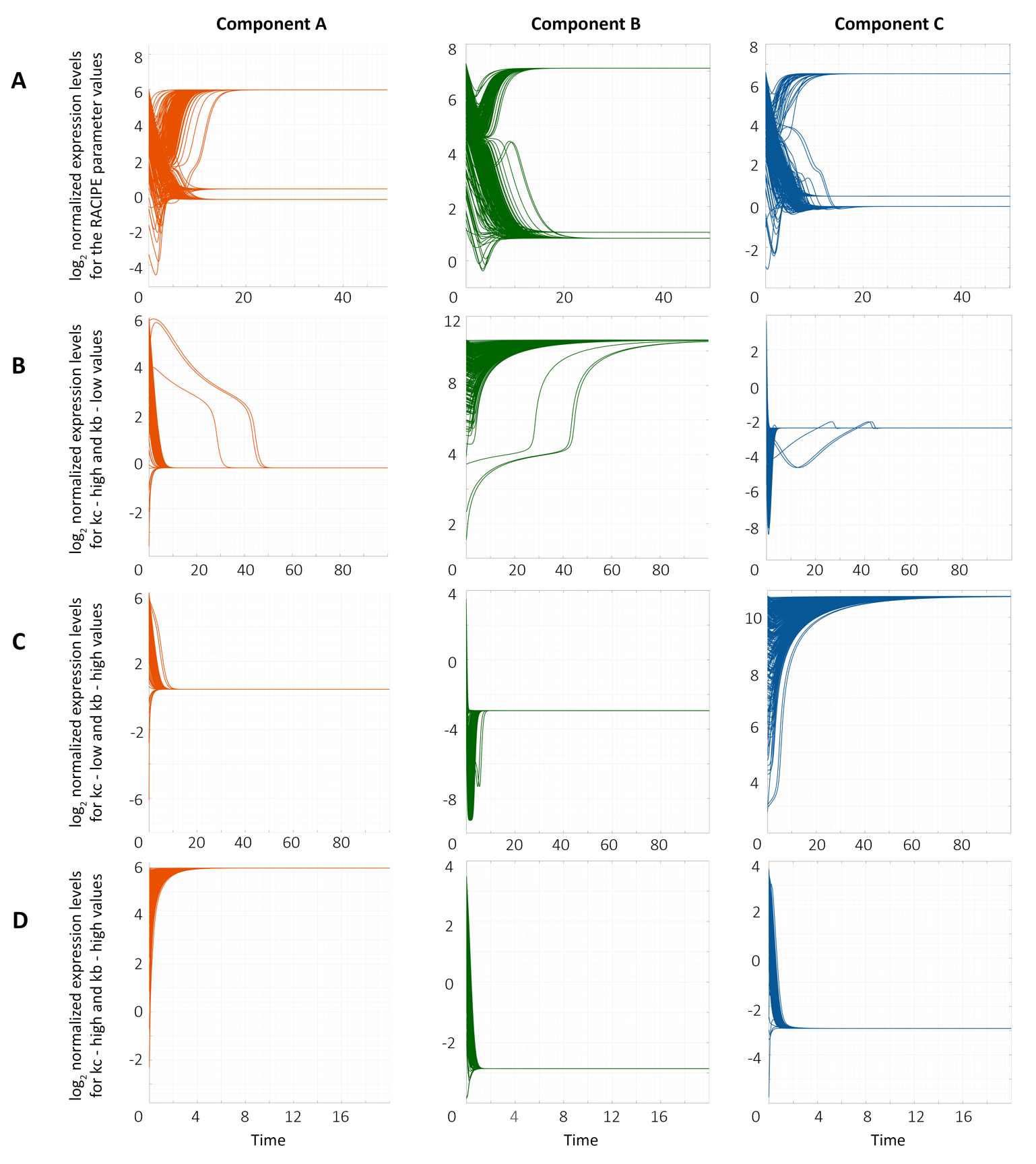

**Figure S12**: Dynamics plot for a representative parameter case of tristable solution of the toggle triad. A) Dynamics plots for the parameter values given by RACIPE, showing (high A, low B, low C), (low A, high B and low C) and (low A, low B, high C) states. B-D) Same as A) but for different kb and kc values. All parameter values are given in Table S13.

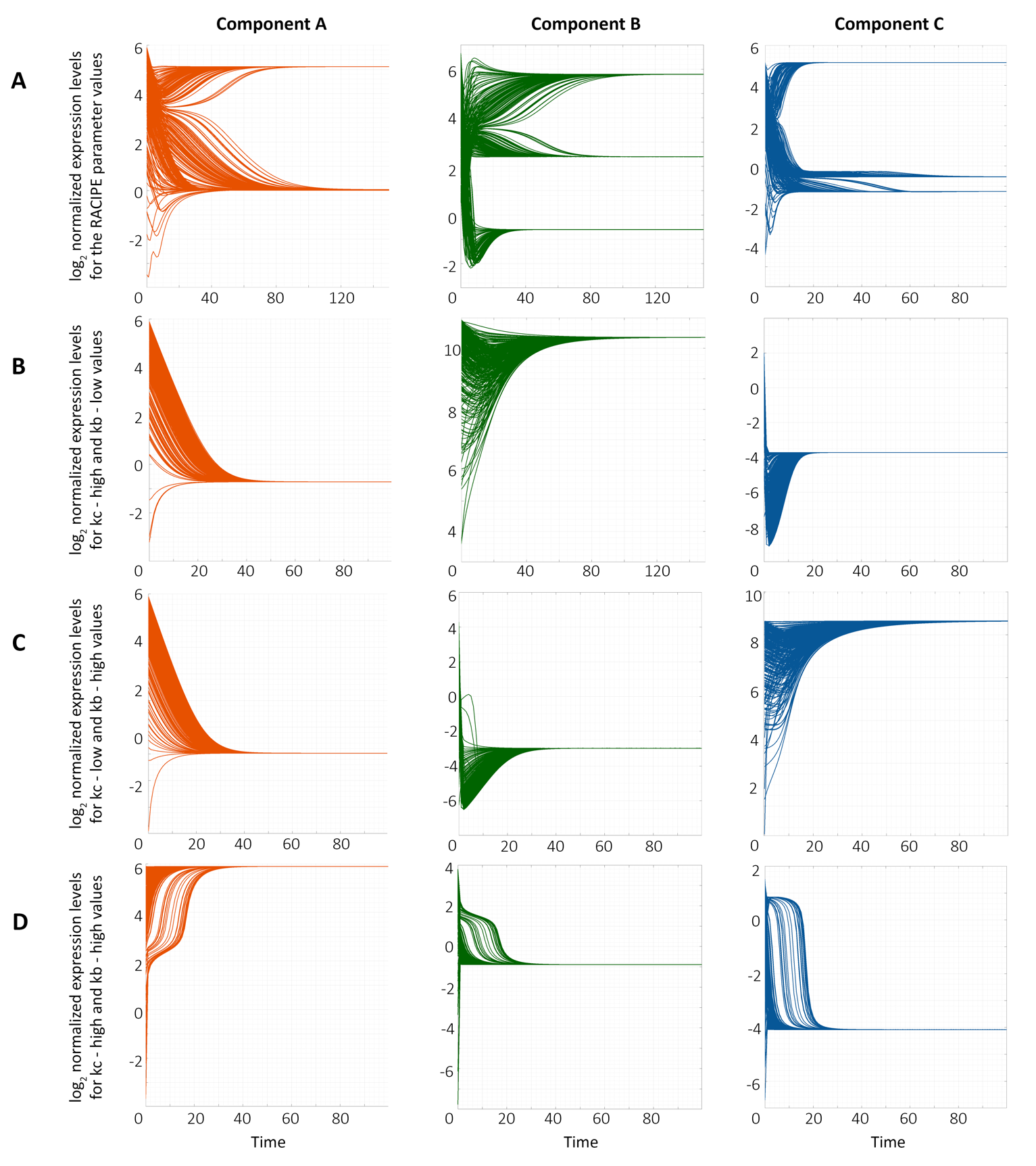

**Figure S13**: Dynamics plot for a representative case of tristable solution of the toggle triad. A) The dynamics plots for the parameter values given by RACIPE, showing (high A, low B, low C), (low A, high B and low C) and (low A, low B, high C) states. B-D) Same as A) but for different values of kb and kc. All parameter values are given in Table S13.

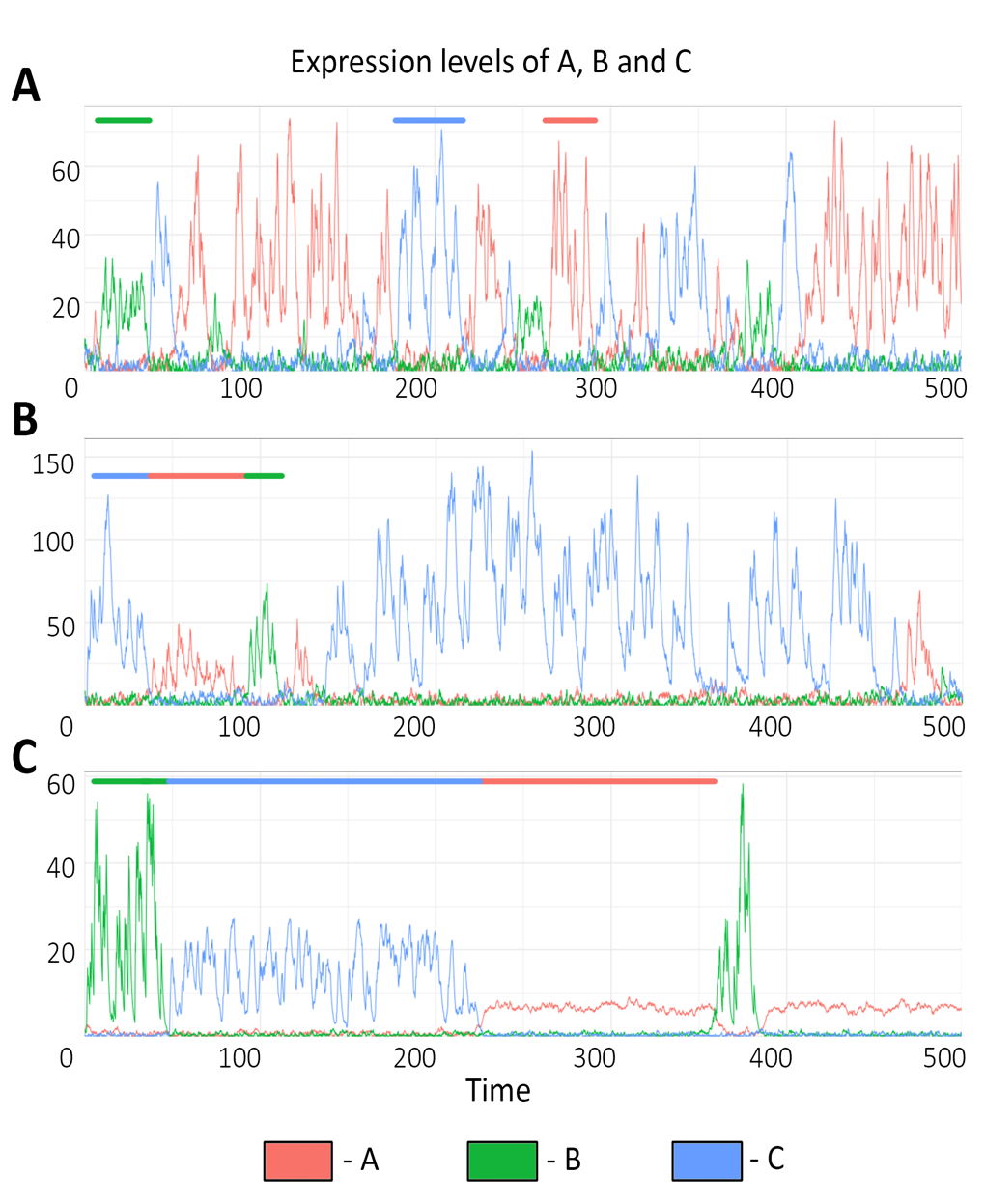

**Figure S14**: sRACIPE results for another replicate of parameter sets used in Fig 7. A) Dynamics plot showing switching between states for parameter set 1. Color bars on top representatively mark the regions of each of the states – green bar shows (low A, high B, low C), red bar shows (high A, low B, low C) and blue bar shows (low A, low B, high C). B, C) Same as A) but for a replicate of parameter set 2 and 3 respectively. Corresponding parameter values are given in Table S13.

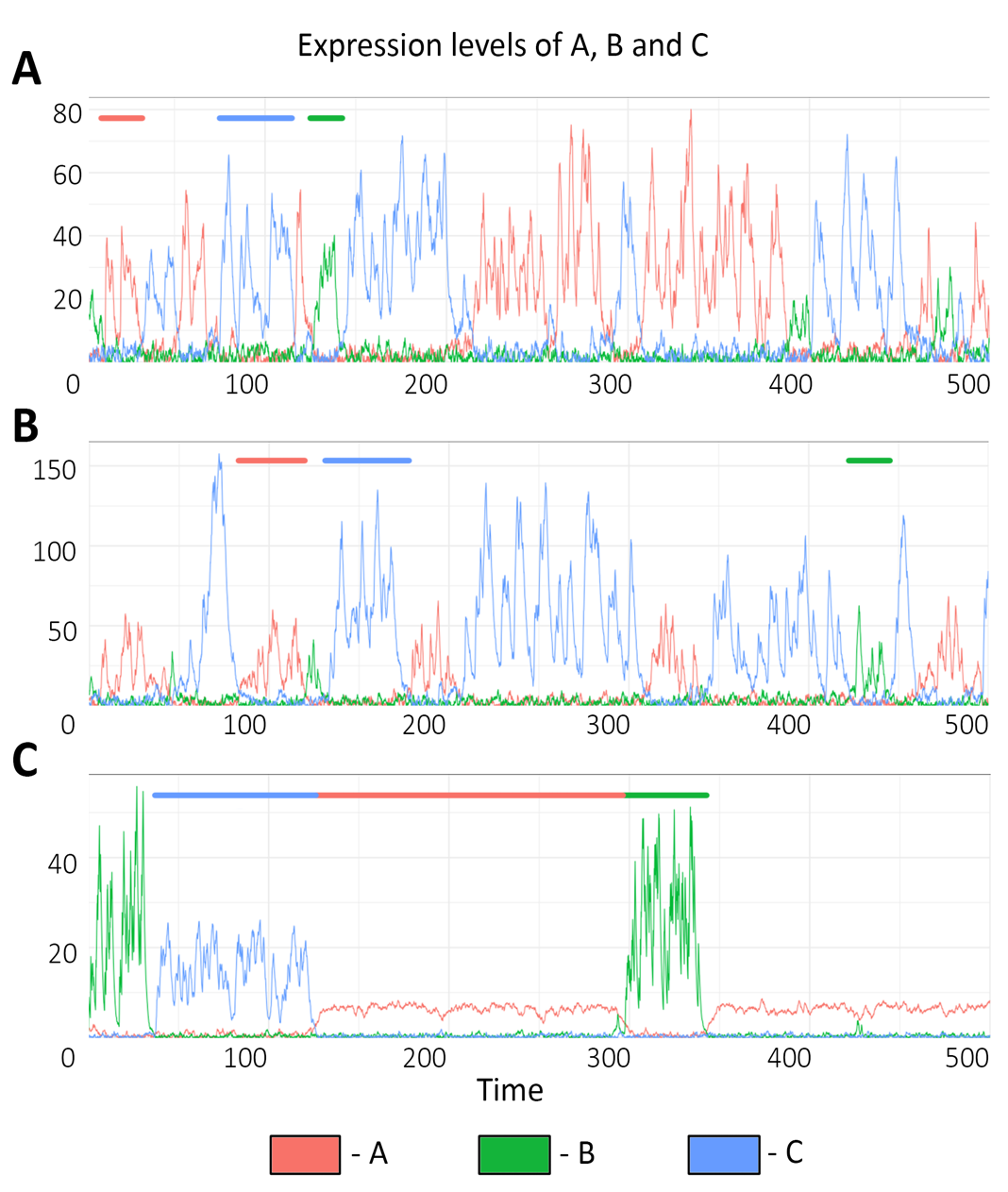

**Figure S15**: sRACIPE results for another replicate of parameter sets used in Fig 7. A) Dynamics plot showing switching between states for parameter set 1. Color bars on top representatively mark the regions of each of the states – green bar shows (low A, high B, low C), red bar shows (high A, low B, low C) and blue bar shows (low A, low B, high C). B, C) Same as A) but for a replicate of parameter set 2 and 3 respectively. Corresponding parameter values are given in Table S13.

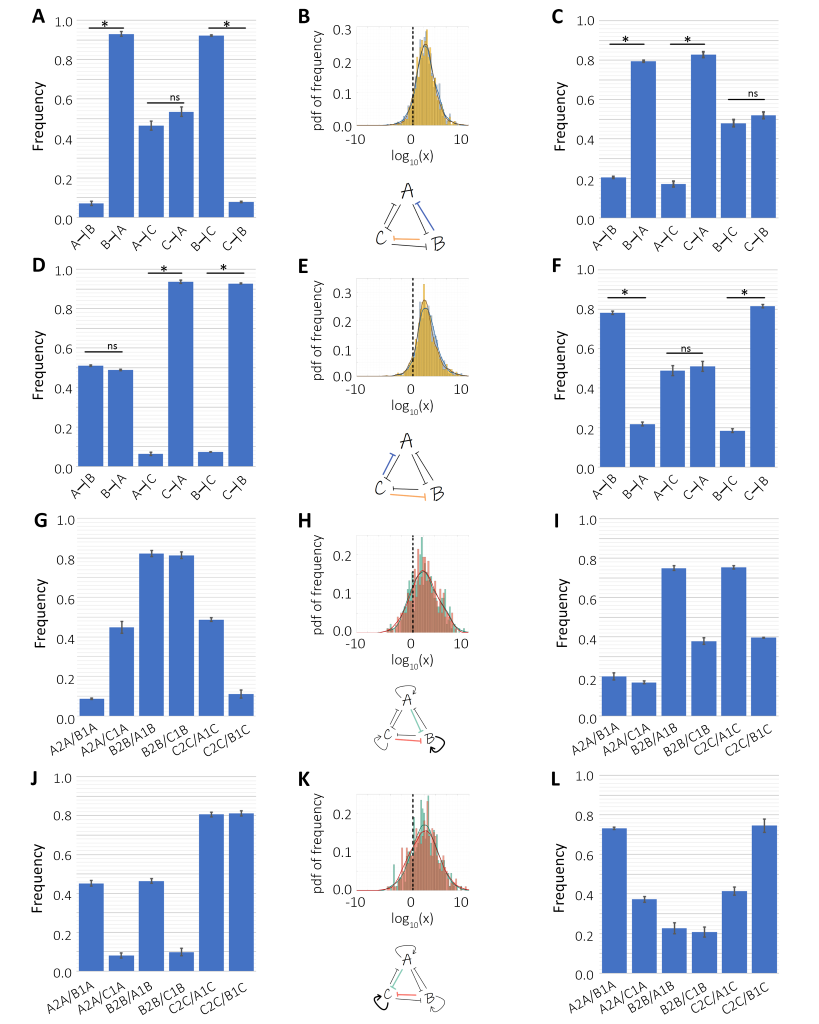

**Figure S16**: Analysis of the strength of interactions between components in monostable and bistable states for toggle triad (TT) and that with self-activation (TT+3SA). A, B) Same as Fig 8A,B but for monostable {aBc} case for TT. Schematic shows that inhibitory links from B to A and C are stronger than inhibition links from A and C to B. Probability distribution functions of histograms of frequency of values of x(B–|A)/x(A–|B) (blue) and values of x(B–|C)/x(C–|B) (yellow) in TT for {Abc}. C) Same as Fig 8C but for bistable phase {abC, aBc} case for TT. D, E) Same as Fig 8 A, B but for monostable {abC} case for TT. Schematic shows that inhibitory links from C to A and B are stronger than inhibition links from A and B to C. Probability distribution functions of histograms of frequency of values of x(C–|A)/x(A–|C) (blue) and values of x(C–|B)/x(B–|C) (yellow) in TT for {abC}. F) Same as Fig 8C but for bistable {abC, Abc} case for TT. G,H) Analysis of monostable {aBc} case for TT+3SA. Probability distribution functions of histogram of frequency of x(B2B)/x(A1B) and x(B2B)/x(C1B) in TT+3SA. The x-axis is log_10_ transformed and the dotted line represents the numerical value of 1. Schematic showing that for most of the parameter sets corresponding to {Abc}, self-activation of B dominates inhibition of B by A or C. I) Analysis of bistable {abC, aBc} case for TT +3SA. J) Analysis of monostable {abC} case for TT +3SA. K) Probability distribution functions of histogram of frequency of (C2C/A1C) and (C2C/B1C) in TT+3SA over a log_10_ transformed x-axis and the dotted line representing 1 in normal scale. L) Analysis of the bistable {abC, Abc} case for TT +3SA.

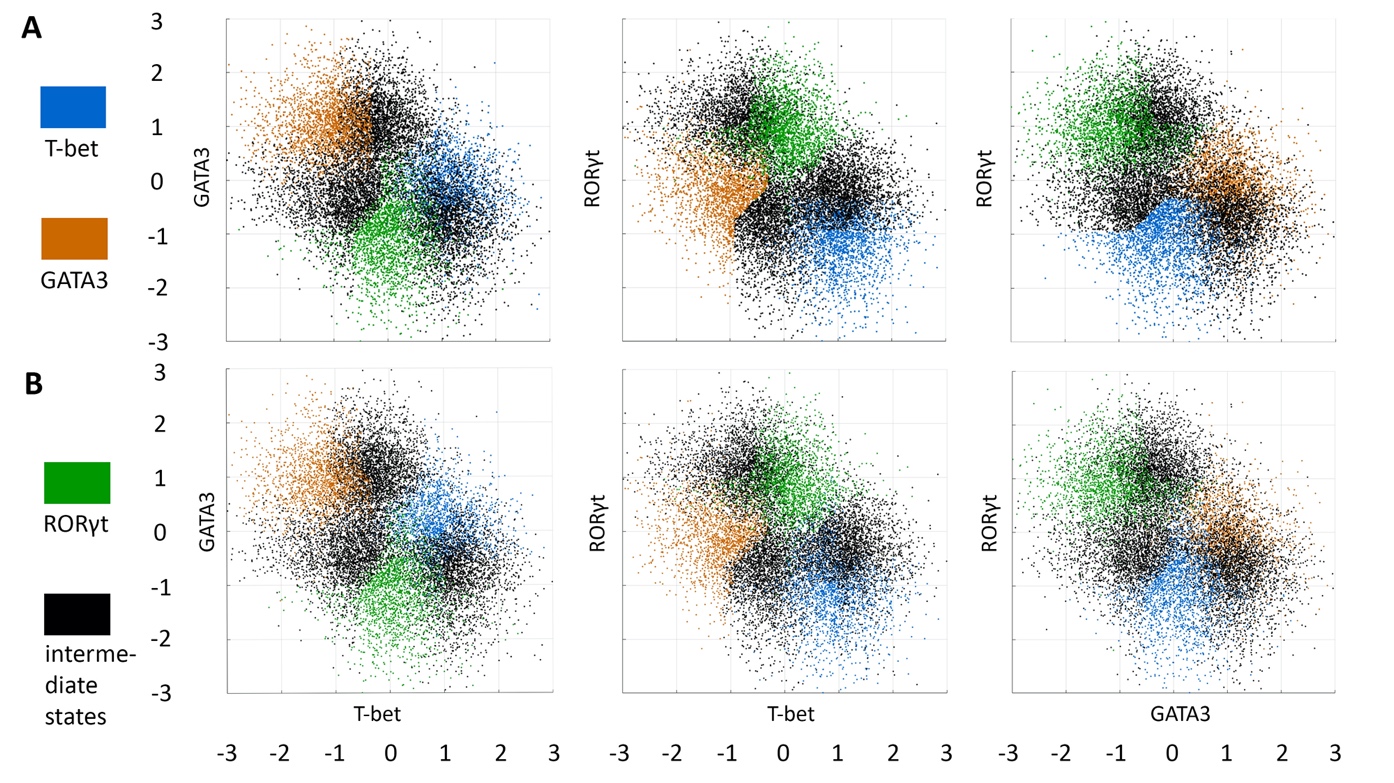

**Figure S17**: K-means clustering for toggle triad (TT). A) The scatter plot obtained for the second replicate of k-means. B) The scatter plot obtained for the second replicate of k-means.

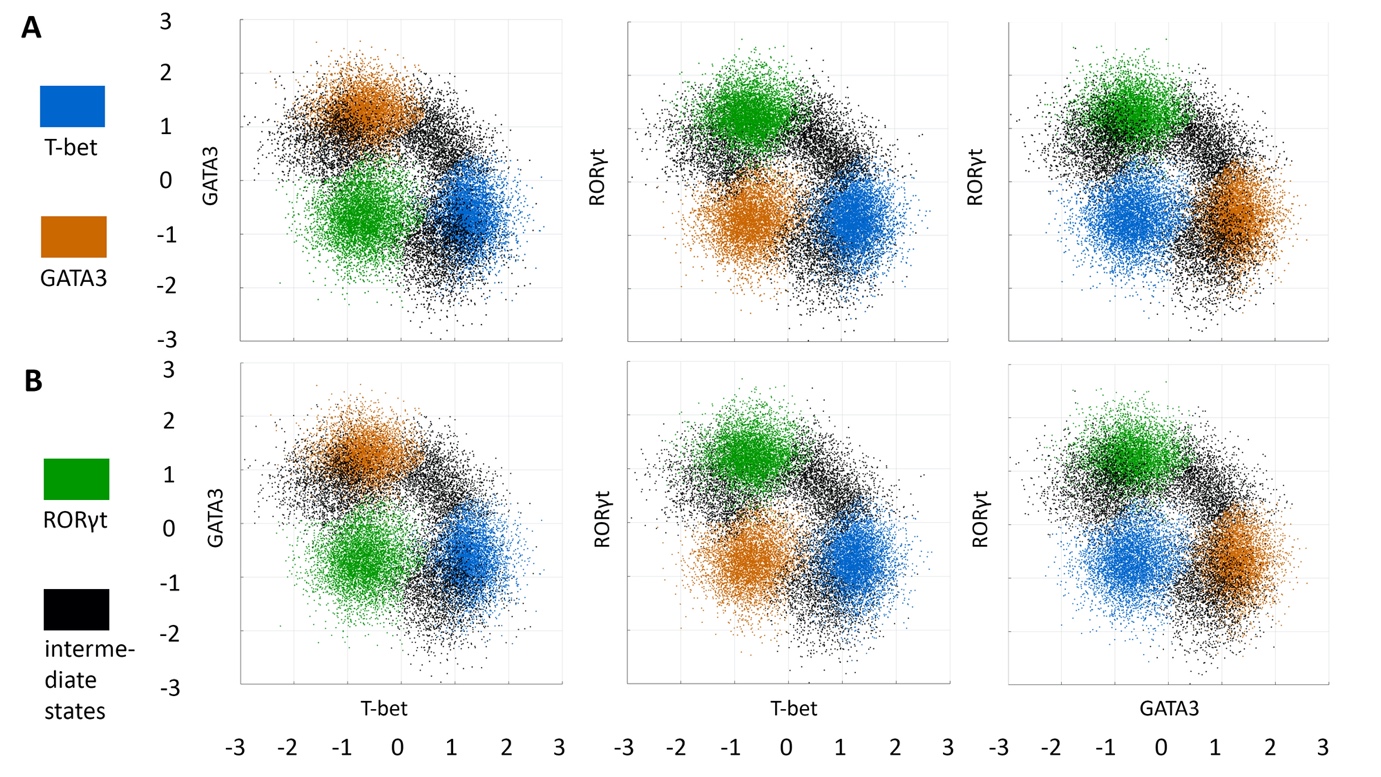

**Figure S18**: K-means clustering for toggle triad with 3 self-activations (TT+3SA). A) The scatter plot obtained for third replicate of k-means. B) The scatter plot obtained for third replicate of k-means.
